# Gene expression inference from cell-free DNA using uncertainty-aware deep learning

**DOI:** 10.64898/2026.02.10.705188

**Authors:** Robert D. Patton, A. Patrick McDeed, Alexander Netzley, Aditya Pawar, Thomas W. Persse, Akira Nair, Patricia C. Galipeau, Ilsa M. Coleman, Pushpa Itagi, Pooja Chandra, Erolcan Sayar, Mohamed Adil, Manasvita Vashisth, Joseph B. Hiatt, Ruth Dumpit, Lori Kollath, Ridvan Arda Demirci, Alireza Ghodsi, Hung-Ming Lam, Colm Morrissey, Delphine L. Chen, Michael T. Schweizer, Amir Iravani, Andrew C. Hsieh, David MacPherson, Michael C. Haffner, Peter S. Nelson, Gavin Ha

## Abstract

Tumor gene expression profiling provides crucial diagnostic information for guiding therapy, but standard tissue biopsies are invasive, spatially biased, and may inadequately sample metastatic disease. Cell-free DNA (cfDNA) provides a minimally invasive alternative for tumor genotyping, yet reconstructing robust, transcriptome-wide expression from standard-depth cfDNA whole-genome sequencing (WGS) remains a major challenge. We developed a deep learning framework comprising Triton, for comprehensive cfDNA feature extraction, and Proteus, a probabilistic model that infers single-gene expression from standard-depth cfDNA WGS. Proteus outperformed prior cfDNA approaches in reconstructing molecular phenotypes from matched tumor transcriptomes across multiple cancer types, including prostate, lung, and bladder cancer cohorts, with uncertainty-guided withholding improving model reliability. Proteus further enabled assessment of therapeutic target activity, prognostic transcriptional programs, and candidate treatment-emergent resistance states, establishing a generalizable framework for minimally invasive functional genomics in precision oncology.

## MAIN

Cancer genomic profiling provides important diagnostic and prognostic information capable of enhancing treatment allocation and improving outcomes. However, clinically actionable tumor states are increasingly defined by their gene expression profiles, which provide complementary information capable of classifying tumor subtypes and identifying therapeutic targets.^1–3^ Tumor biopsies are the common substrate for genomic, transcriptomic and immunohistochemical assays, but they are invasive procedures associated with high costs and morbidity and are often difficult to acquire in patients with advanced cancers.^4^ A simple blood draw, as a minimally invasive “liquid biopsy,” provides a promising alternative, but inferring gene expression directly from plasma remains challenging.^5,6^

Recent liquid biopsy approaches have mapped cell-free DNA (**cfDNA**) fragmentation patterns, characterized methylation and histone modifications, or quantified cell-free RNA to identify cancer-associated transcriptional activity and phenotypic signatures.^7,8^ However, many rely on specialized epigenomic or immunoprecipitation-based assays (e.g., APEX), require patient-specific customized assays, or employ ultra-deep targeted sequencing (e.g., panel-optimized EPIC-Seq), limiting their scalability and broad adoption.^9–12^ Standard-depth whole-genome sequencing (**WGS**) of liquid biopsy-derived cfDNA offers a more accessible alternative. Because cfDNA in circulation is primarily composed of nucleosome-protected fragments, its coverage and fragmentation patterns retain information about the chromatin states of the contributing cells.^13–15^ WGS of cfDNA can capture these features, but translating them into accurate and reliable estimates of single-gene expression in RNA-seq units remains a computational challenge.^16–18^ Furthermore, deterministic methods that force an expression estimate even when the underlying tumor signal is weak may produce misleading and potentially harmful results. A clinically interpretable framework should therefore explicitly quantify uncertainty, ensuring that ambiguous predictions can be withheld or down-weighted rather than overinterpreted.

We hypothesized that integrating fragmentation, nucleosome-positioning and sequence features from standard-depth cfDNA WGS would enable robust gene expression inference. We therefore developed Triton, for comprehensive cfDNA feature extraction, and Proteus, which integrates these features with promoter sequence and gene annotations while directly modeling tumor- and background-associated cfDNA contributions. Proteus predicts a distribution over RNA-seq-like expression values for each gene–sample pair, enabling uncertainty-aware downstream analyses. We developed and evaluated the framework using patient-derived xenograft (**PDX**) circulating tumor DNA (**ctDNA**), healthy-human donor cfDNA, controlled *in silico* admixtures and patient cohorts with matched tumor RNA-seq across castration-resistant prostate cancer (**CRPC**), small-cell lung cancer (**SCLC**) and bladder cancer (**BLCA**). We then benchmarked the recovery of gene-level expression variation against simpler models and a prior cfDNA approach, evaluated generalization in held-out patient cohorts, and tested whether uncertainty-guided abstention improved the reliability of individual gene-activity calls. Finally, we applied Proteus to clinical cohorts to classify tumor lineage, monitor treatment-associated transcriptional changes and assess clinically relevant therapeutic target activity.

## RESULTS

### Triton captures transcription-associated cfDNA features at base-pair resolution

Cell-free DNA fragmentation patterns reflect DNA protection from nuclease digestion, indicating nucleosome occupancy and chromatin organization.^13,19^ Actively transcribed genes exhibit a nucleosome-depleted region (**NDR**), a stably located +1 nucleosome and increased promoter and gene-body fragment-length heterogeneity, whereas inactive genes show regular nucleosome positioning and less fragment-length diversity (**Fig. 1a,b**).^19–21^ To capture these patterns, we expanded our previously reported method Triton,^16^ which extracts coverage, fragmentation and phased-nucleosome features from cfDNA WGS, producing both region-level features and spatial signals at base-pair resolution (**Fig. 1a–c; Supplementary Fig. 1a,b; Methods**). Triton resolves transcription-associated fragmentation patterns across both promoters (transcription start site [TSS] ±1,000 bp) and gene bodies while incorporating a fragment-length-based correction to improve nucleosome localization, extending prior promoter-centric approaches.^10,11,14^ As a standalone tool, Triton can also profile composite, user-defined open-chromatin regions and transcription factor binding sites, supporting diverse applications including transcription factor activity inference, molecular phenotyping and, here, transcriptome reconstruction using Proteus.^16^

**Figure 1.**
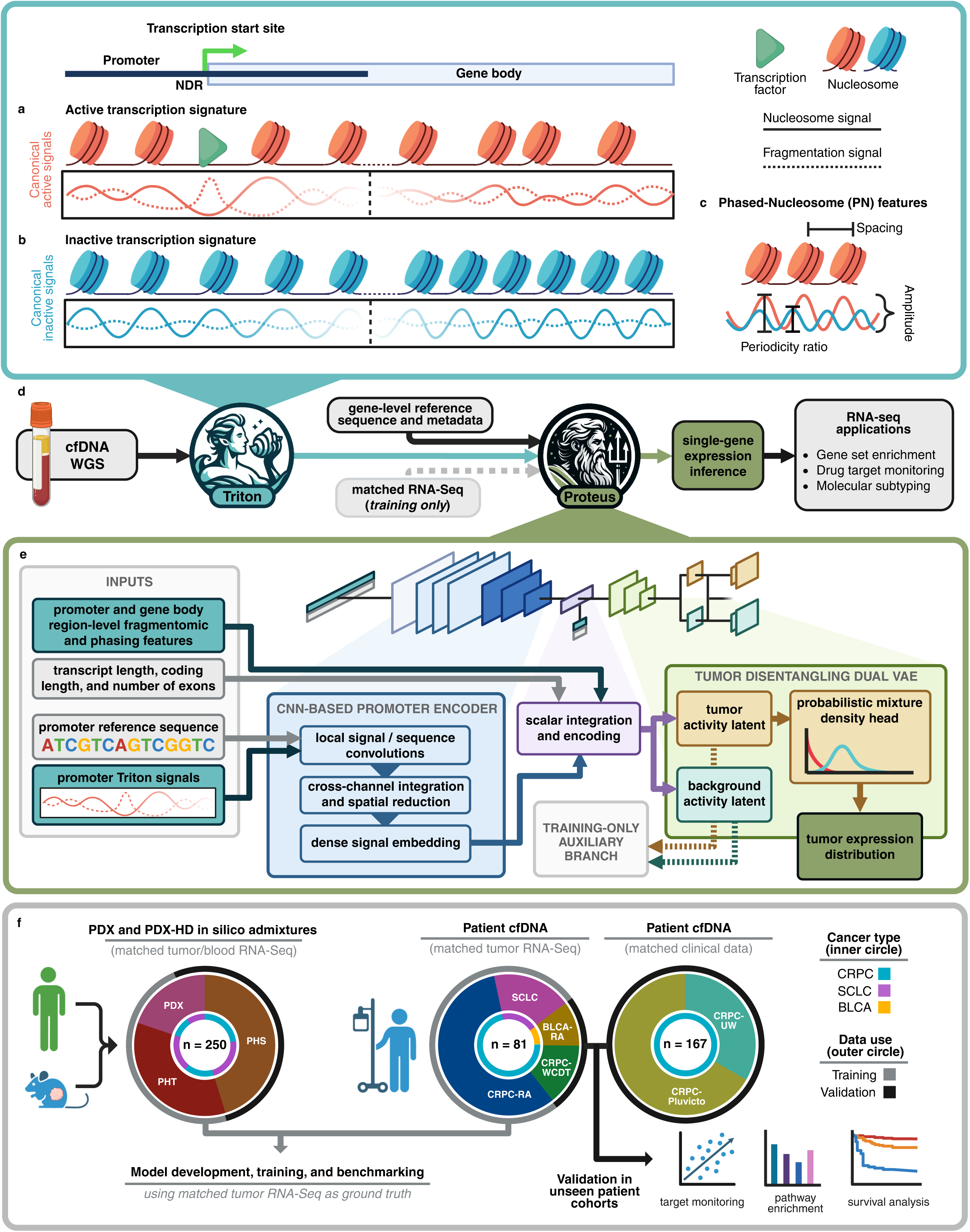
Proteus framework for uncertainty-aware tumor gene expression inference from cell-free DNA. **a)** Canonical nucleosome coverage and cfDNA fragmentation signals in the TSS region (left) and body (right) of a gene undergoing active transcription. In the promoter, binding of TFs at the TSS leads to a nucleosome-depleted region (NDR), or dip in coverage, while the regulatory role of the +1 nucleosome leads to more stable positioning, seen as a peak in coverage. Accessible promoter chromatin and heterogeneous cfDNA cleavage generate fragments of diverse lengths that inversely mirror nucleosomal protection, with higher diversity expected at the NDR. In the body of the gene, active transcription is associated with less consistent positioning, reflecting chromatin remodeling, which also leads to greater heterogeneity in fragmentation patterns. **b)** In the case of an inactive gene, no NDR or highly phased +1 peak is expected at the TSS, and regular nucleosome positioning extends into the gene body reflecting inactive chromatin. More consistent fragmentation patterns, typical of nucleosome protection, are found in both regions. **c)** From the phased-nucleosome (PN) signal produced by Triton, three additional region-level metrics capturing chromatin structure are extracted: PN spacing, which reveals the mean inferred distance between nucleosomes; PN amplitude, which measures the strength of well-phased nucleosome signal components; and PN periodicity ratio, which reports the relative contribution of longer versus shorter nucleosome periodicities within the signal. **d)** Overall analytical workflow. Triton nucleosome and fragmentation spatial signal profiling from cfDNA WGS at individual gene TSS regions, as well as region-level features extracted from both TSS and gene bodies (a–c), are combined with matched metadata (transcript length, coding region length, number of exons, and TSS region reference sequence) as inputs to Proteus. Matched RNA-seq of tumor biopsies is used for model training and evaluation. As a final output, Proteus predicts individual tumor gene expression as probability distributions of RNA-seq log_2_(CPM+1) abundance, which can be used in uncertainty-aware downstream analyses including gene set enrichment, therapeutic target monitoring, and molecular subtyping. **e)** Proteus model architecture. Inputs include TSS-centered Triton signal profiles, promoter reference sequence, promoter and gene-body Triton scalar features and gene-level reference annotations. A convolutional promoter encoder learns local sequence and cfDNA signal representations, which are integrated with scalar features and passed to dual variational encoders representing tumor-associated and background-associated activity. An auxiliary TFx predictor and learned weight predictor condition recombination of tumor and background latents for internal reconstruction, while tumor and background mixture-density heads output hurdle–gamma–normal expression distributions. Point estimates are reported as mixture means and prediction intervals are derived from mixture quantiles. **f)** Study design and cohort overview. Proteus was developed, trained, and benchmarked using patient-derived xenograft and healthy-donor in silico admixtures with matched tumor RNA-seq and available healthy-donor blood RNA-seq (left) along with patient cfDNA with matched tumor biopsy RNA-seq (center). The framework was evaluated in both patient cfDNA cohorts with matched tumor RNA-seq and in clinical cfDNA cohorts without contemporaneous matched tumor RNA-seq. Inner circles denote cancer type, middle circles denote cohort, and outer circles denote data use. CRPC, castration-resistant prostate cancer; SCLC, small-cell lung cancer; BLCA, bladder cancer; HD, healthy donor; PDX, patient-derived xenograft; TFx, tumor fraction; WGS, whole-genome sequencing.

### Development of a deep learning framework for gene expression inference from cfDNA

To predict tumor-associated transcriptional activity at individual gene loci from cfDNA, we then developed a multimodal deep learning framework termed Proteus that integrates Triton features with gene promoter reference sequences and annotations (**Fig. 1d; Methods**).^22–24^ A convolutional neural network jointly encodes promoter spatial features and reference sequence before integrating promoter and gene-body regional features (**Fig. 1e**). Dual variational autoencoders model tumor-associated and background cfDNA contributions, while tumor fraction (**TFx**) provides auxiliary supervision only during training and is not supplied at inference. Finally, to derive point predictions and model-based uncertainty measures, a mixture density network outputs a probability distribution over log_2_(CPM+1) expression values for each gene-sample pair (**Supplementary Fig. 1c; Methods**). Genes are processed individually without providing gene/sample identity or tissue/cancer-type labels^25^ to discourage overfitting and focus predictions on local cfDNA features, limited sequence context and transcript metadata.

We employed a staged study design that separated training, benchmarking, matched-tissue validation, and clinical application (**Fig. 1f; Methods**). Model development used pure ctDNA derived from 40 PDX models (20 CRPC and 20 SCLC), healthy donor cfDNA (HD-L, n = 10), their in silico admixtures (n = 80), and two patient cohorts with matched tumor RNA-seq (CRPC-RA, n = 44; SCLC^10^, n = 16). Ten additional PDX models (5 CRPC and 5 SCLC) and ten healthy donor samples (HD-U) were excluded from all model development and used to create controlled benchmarking admixtures (n = 120). Externally generated CRPC-WCDT^26^ (n = 13) and BLCA-RA (n = 8) cohorts with paired cfDNA and RNA-seq from tumor tissue biopsies were used for real-world benchmarking and to assess generalization. Finally, we further evaluated Proteus in two metastatic CRPC cohorts without matched RNA-seq (n = 167**, Supplementary Table 1**) to illustrate transcriptional analysis applications in precision oncology.

### Gene-specific correlations distinguish informative cfDNA expression predictions

Prior plasma-based expression-inference studies have evaluated accuracy using correlations calculated across individual genes or aggregated gene groups within each sample.^7,10^ However, because the ordering of highly and lowly expressed genes is largely conserved across samples derived from the same organ, tissue or cell type, and across tumors exhibiting similar phenotypes, sample-level concordance can remain high even for biologically uninformative predictions. To illustrate this distinction, we compared RNA-seq-only controls assigning each gene its expression in a random alternative sample (“naïve predictor”) or assigning the entire profile of the closest transcriptional neighbor (**“**nearest-neighbor predictor**”; Methods**). Both retained high sample-level correlation (mean ρ_sample_ = 0.91 and 0.87 for nearest-neighbor and naïve predictors), but only the nearest-neighbor control recovered gene-specific variation across samples (mean ρ_gene_ = 0.36 versus 0.00; **Fig. 2a**). To measure performance at the cohort level, we therefore adopted a gene-variance-weighted correlation (ρ_GVW_), which emphasizes genes with greater observed variance between patients (**Methods**).

**Figure 2.**
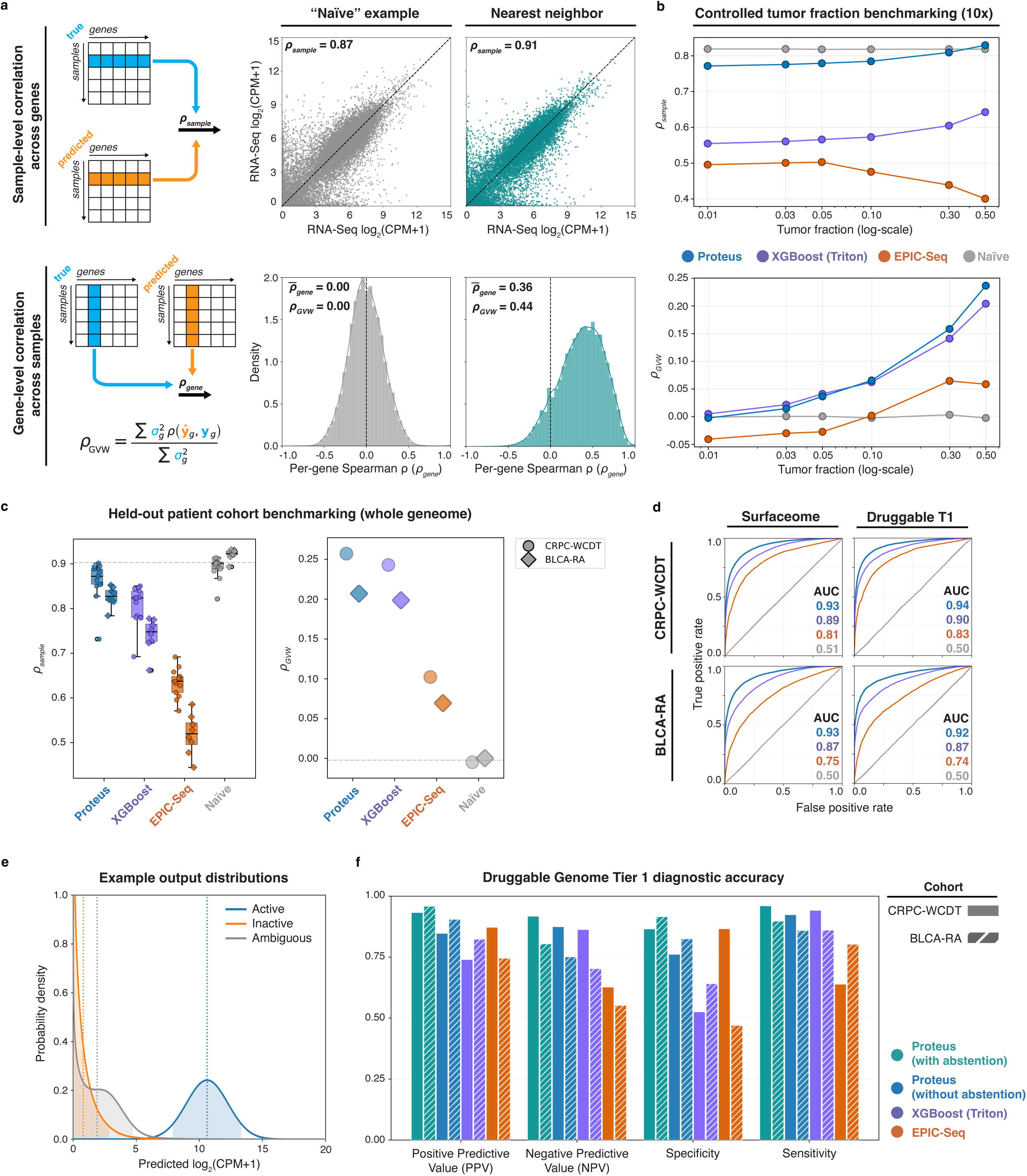
Benchmarking Proteus for single-gene expression inference and uncertainty-aware activity calls. **a)** Overview of benchmarking metrics used to distinguish global sample-level agreement from gene-specific expression recovery. Sample-level correlation across single genes can remain high even for uninformative predictions because most samples share a broadly similar transcriptome-wide abundance distribution (top row). Gene-level Spearman correlations across samples and gene-variance-weighted correlation (ρ_GVW_, bottom row) were therefore used to evaluate whether predictions recover sample-specific variation for individual genes. In the “naïve” example (left), sample-level Spearman correlation remains high despite per-gene correlations across samples being centered at zero. In the nearest-neighbor example (right), sample-level Spearman correlation is comparable, but there is a distinct, positive shift in gene-level correlations. **b)** Controlled tumor-fraction benchmarking at 10× WGS depth across tumor fractions from 0.01 to 0.50, with n = 10 held-out PDX–donor admixtures per TFx/depth condition. Proteus was compared with an XGBoost model trained on Triton features, EPIC-Seq in WGS mode and a naïve baseline. Using sample-level Spearman correlation across single genes (ρ_sample_, top) reveals only a weak dependence on tumor fraction, reflecting that cross-gene correlation will tend to overestimate predictive accuracy as illustrated in (a), whereas gene-variance-weighted Spearman correlation (ρ_GVW_, bottom) taken across samples shows a clear dependence on TFx. **c)** Whole-genome benchmarking in held-out patient cfDNA cohorts with matched tumor RNA-seq. Per-sample (across single genes) Spearman correlations are shown for individual samples (left), and ρ_GVW_ is shown at the cohort level (right) for CRPC-WCDT (n = 13) and BLCA-RA (n = 8); cohort is indicated by marker shape: circles for CRPC-WCDT and diamonds for BLCA-RA. **d)** Receiver-operating characteristic curves for on/off gene-expression classification in Cancer Surfaceome and Druggable Genome Tier 1 gene sets, evaluated in CRPC-WCDT and BLCA-RA. AUC values are shown for Proteus, XGBoost/Triton, EPIC-Seq and the naïve baseline and are indicated by color. Cancer Surfaceome: BLCA-RA n = 26,546 gene–sample pairs; CRPC-WCDT n = 43,252 gene–sample pairs. Druggable Genome Tier 1: BLCA-RA n = 11,027 gene–sample pairs; CRPC-WCDT n = 17,941 gene–sample pairs. **e)** Example Proteus-predicted *AR* expression distributions for CRPC-RA samples classified as active (17-113; point estimate = 10.6, AOR = 10.8), inactive (18-039; point estimate = 0.8, AOR = -7.2) or ambiguous (05-144; point estimate = 1.91, AOR = -0.3). Predictions were classified as active at AOR > 5, inactive at AOR < −5 and ambiguous at |AOR| < 5. Curves show predicted expression distributions, shaded regions show central 90% predictive intervals and vertical dotted lines show point estimates. **f)** Diagnostic accuracy of Druggable Genome Tier 1 on/off calls, including positive predictive value, negative predictive value, specificity and sensitivity, for Proteus with and without uncertainty-informed abstention, XGBoost/Triton and EPIC-Seq. Solid bars denote CRPC-WCDT and hatched bars denote BLCA-RA. Uncertainty-informed Proteus calls used |AOR| < 5 as the default ambiguous-call threshold; abstained gene-sample pairs were excluded from PPV, NPV, specificity and sensitivity calculations.

We benchmarked Proteus in controlled *in silico* admixtures generated at 5× and 10× WGS depth across tumor fractions from 1% to 50% to test performance in low-depth and low-TFx settings (n = 10 per TFx/depth condition, using unseen data; **Methods**). Comparators included an XGBoost model trained directly on Triton features, representing a simpler machine learning approach, EPIC-Seq^11^ applied in WGS mode (as opposed to its optimized targeted assay), and the naïve control (**Fig. 2b; Methods**). As expected, sample-level correlation remained high in the 10× benchmarking even for the naïve control, and ρ_sample_ showed only modest dependence on TFx across all tested models. By contrast, ρ_GVW_ revealed a strong TFx dependence and clearly separated models by their ability to recover gene-level expression variation. Proteus performed best across the admixture series, whereas XGBoost provided reduced performance and EPIC-Seq showed weaker gene-level recovery under these conditions. The same pattern with reduced overall accuracy was observed at 5× depth (**Supplementary Fig. 2a**). Thus, gene-expression prediction is strongly TFx dependent, and Proteus more effectively converts standard-depth cfDNA WGS features into gene-expression estimates than either a simpler machine learning model or EPIC-Seq under matched WGS conditions.

### Integrated cfDNA features predict tumor expression in unseen cohorts

We next evaluated Proteus in two complementary patient cfDNA cohorts with matched tumor RNA-seq that were excluded from model development: externally generated CRPC-WCDT^26^ (*n* = 13, depth = 95–150×, TFx = 0.19–0.66) and BLCA-RA (*n* = 8, depth = 55–94×, TFx = 0.08–0.85), representing an unseen cancer type. Proteus achieved the strongest combined performance across both cohorts, maintaining high sample-level and gene-variance-weighted correlations (CRPC-WCDT [mean ρ_sample_, ρ_GVW_] = [0.86, 0.26]; BLCA-RA = [0.83, 0.21]; **Fig. 2c; Supplementary Table 2**). XGBoost trained on Triton features showed moderately reduced performance (CRPC-WCDT = [0.81, 0.24]; BLCA-RA = [0.74, 0.20]), while EPIC-Seq in WGS mode was weaker across both cohorts and metrics (CRPC-WCDT = [0.63, 0.10]; BLCA-RA = [0.52, 0.07]). The EPIC-Seq sample-level correlations were consistent with the median correlation reported for an independent WGS implementation in the APEX study (ρ_sample_ = 0.61), in which APEX itself achieved ρ_sample_ = 0.82 using cfChIP-seq.^12^ As anticipated, the naïve predictor retained high apparent sample-level concordance but no gene-level expression recovery (CRPC-WCDT = [0.89, 0.00]; BLCA-RA = [0.92, 0.00]). A sequence-only version of Proteus similarly failed to recover gene-level expression variation, whereas omitting promoter sequence reduced transcriptome-wide accuracy (**Supplementary Fig. 2b**). We also benchmarked binary classification of expression activity using the Cancer Surfaceome^27^ and Druggable Genome Tier 1^28^ gene sets (**Methods**). Proteus achieved the highest AUROC in both held-out cohorts (0.92– 0.94), followed by XGBoost (0.87–0.90), EPIC-Seq (0.74–0.83) and the naïve control (0.50–0.51; **Fig. 2d**). Together, these results support integrating promoter sequence with spatial cfDNA signals to improve gene-expression prediction across independent cohorts and cancer types.

### Uncertainty-aware cfDNA-based predictions improve gene-activity call reliability

A distinguishing feature of Proteus is that it models each gene’s expression as a probability distribution comprising an active “on” component and lower-expression or inactive “off” components (**Methods**). For each gene–sample pair, Proteus reports both a point estimate and a 90% predictive interval (**PI**), enabling direct assessment of prediction uncertainty. We further summarized confidence in binary gene activity calls using the activity odds ratio (**AOR**), defined as the log₂ odds of the active component relative to the inactive and ambiguous components, enabling model-based prediction abstention using a default threshold of |AOR| < 5 (**Fig. 2e; Supplementary Fig. 3c; Methods**). Both AOR and PI width were associated with absolute prediction error in unseen cohorts, supporting their utility as complementary measures of model uncertainty (**Supplementary Fig. 3a,b**).

In Druggable Genome Tier 1 benchmarking, AOR-based abstention improved positive predictive value (**PPV**), negative predictive value (**NPV**), specificity and sensitivity across CRPC-WCDT and BLCA-RA (**Fig. 2f**), with similar Cancer Surfaceome results (**Supplementary Fig. 3d; Supplementary Table 2**). Across four cohort–gene-set combinations, withholding 24% of calls improved PPV by 8%, NPV by 5%, sensitivity by 4% and specificity by 9%. Frequently withheld genes had lower expression and were closer to the RNA-seq activity boundary, indicating that their underlying activity states were less distinct, and were also enriched for immune-related pathways and nominally associated with neutrophil and hepatocyte signatures (**Supplementary Fig. 4a–d; Supplementary Table 2**). These patterns are consistent with increased difficulty separating tumor and background contributions for active genes in healthy cell populations that contribute heavily to plasma cfDNA. Together, these results demonstrate a practical advantage of probabilistic inference beyond point-estimate accuracy, as Proteus AOR-based abstention can restrict gene activity calls to those predictions supported by sufficient cfDNA evidence.

### cfDNA fragmentation and nucleosome positioning encode transcriptional activity

We used SHAP analysis^29^ to determine whether Proteus expression predictions were supported by biologically interpretable cfDNA features (**Fig. 3a–c; Methods**). Greater fragment length diversity in both the promoter and gene body was associated with higher expression, whereas lower promoter fragment depth was associated with higher expression, consistent with nucleosome depletion at the TSS (**Fig. 3b**). By contrast, gene body features produced more broadly distributed effects, and showed the largest aggregate attributions, extending previous studies that largely focused on the promoter and TSS (**Supplementary Fig. 5**).^10,11,14^ Phased-nucleosome features unique to Triton contributed similarly to gene-body fragmentation features, with increased spacing and periodicity ratio generally associated with higher expression, whereas greater phased-nucleosome amplitude was associated with lower expression. In bp-resolution signals at the promoter, attributions were concentrated near the TSS, with lower NDR and higher +1 nucleosome signals associated with increased expression (**Fig. 3c; Supplementary Fig. 6a,b; Methods**). Fragment-end coverage showed complementary signed effects across these regions, indicating that Proteus learned to integrate local nucleosome positioning and fragment-end enrichment with broader gene-body features.

**Figure 3.**
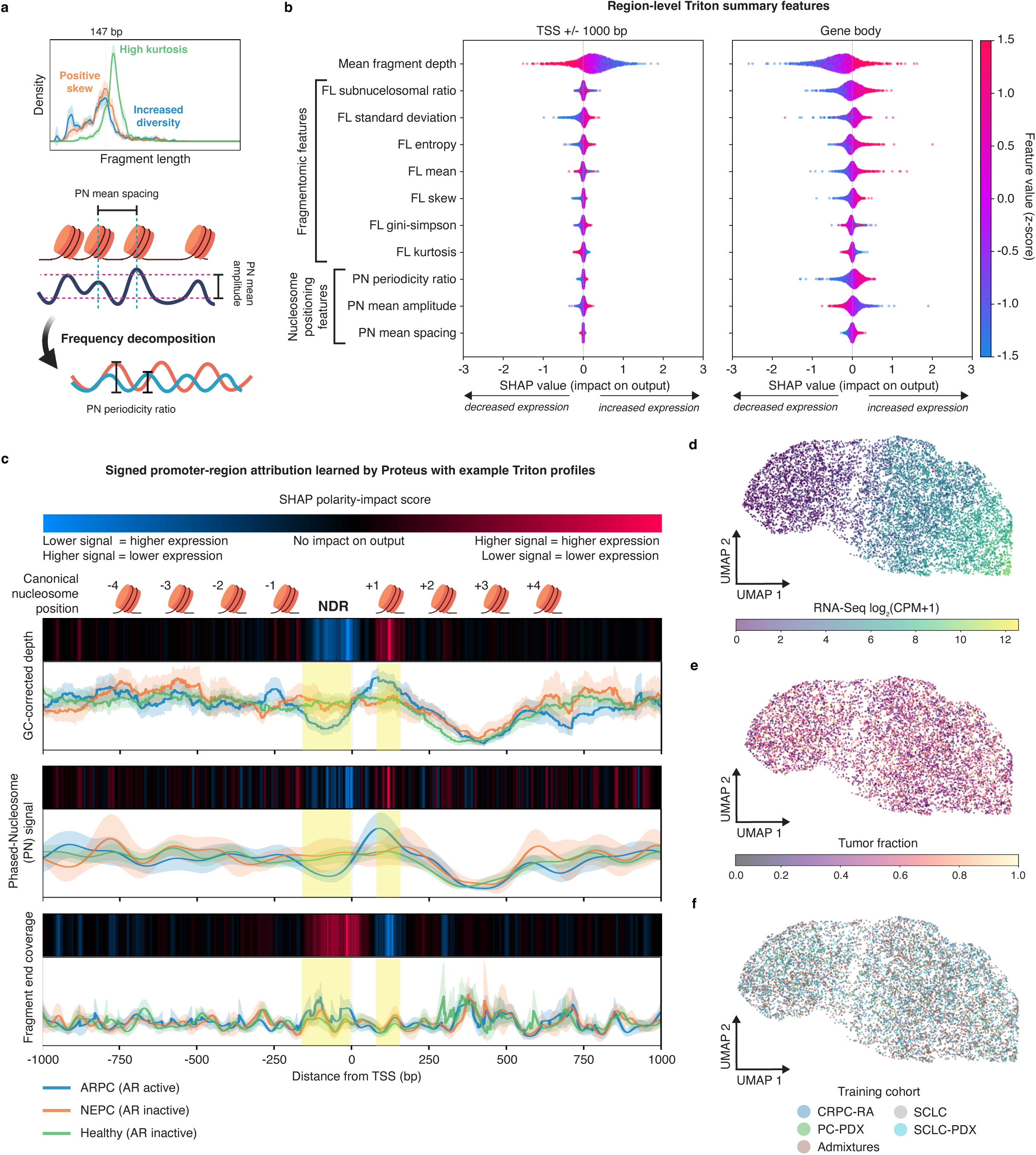
Gene-body fragmentation and promoter nucleosome positioning inform tumor gene expression. **a)** Schematic of Triton region-level feature classes. Fragment-length (FL) features summarize distributional properties of cfDNA fragments, including short-fragment enrichment, mean fragment length, skew, kurtosis and fragment-length diversity. Phased-nucleosome (PN) features summarize inferred nucleosome organization, including mean nucleosome spacing, PN amplitude and a phased periodicity ratio score derived from frequency decomposition of the PN signal. **b)** SHAP values for region-level Triton features used by Proteus, shown separately for features extracted from the TSS ±1,000-bp region and the full gene body (**Methods**). Each point represents a single gene-sample prediction. Point color indicates the feature value after within-sample standardization, except mean fragment depth, which was log-scaled. Position along the x-axis indicates the impact of each feature on predicted expression in log_2_(CPM+1) units. **c)** Signed promoter-region SHAP polarity-impact scores for bp-resolution Triton input signals, aligned to canonical nucleosome positions around the TSS. Red indicates regions where higher signal is associated with higher predicted expression; blue indicates regions where lower signal is associated with higher predicted expression and black indicates limited impact on model output. Example Triton profiles are shown for the AR promoter region in LuCaP PDX ctDNA from ARPC tumors with active AR expression, NEPC tumors with inactive AR expression and healthy donor cfDNA; lines indicate mean signal; shaded bands indicate 95% confidence interval. Yellow shading denotes the nucleosome-depleted region (NDR) and +1 nucleosome region. **d–f)** UMAP projection of the Proteus tumor-activity latent representation colored by matched RNA-seq expression, tumor fraction and training cohort. Each panel shows the same example gene set with different coloration.

UMAP projection^30^ of the tumor-activity latent embeddings further revealed a primary axis corresponding to matched RNA-seq expression, indicating that Proteus organized gene-sample representations by their underlying transcript abundance (**Fig. 3d**). By contrast, the latent embeddings showed no obvious separation by TFx or tissue/cancer type (**Fig. 3e,f**). Model uncertainty also aligned with the activity axis, with AOR, 90% PI width, and predictive error indicating lower confidence and greater error near the on/off embedding boundary (**Supplementary Fig. 7a–c**). Together, these findings demonstrate that Proteus learned established cfDNA signatures of transcriptional activity and organized its latent representations by expression rather than technical or cohort-specific variation.

### Inferred transcriptomes capture tumor phenotypes, pathways, and target expression

We then asked whether Proteus-derived cfDNA expression profiles accurately reproduce tumor-derived whole transcriptome analyses to identify molecular phenotype and pathway activity. Proteus predictions were concordant with tumor RNA-seq at the sample level across CRPC-RA (n = 42 with TFx > 0.05, mean r = 0.90), CRPC-WCDT (n = 13, r = 0.87), BLCA-RA (n = 8, r = 0.82) and SCLC (n = 16, r = 0.88; **Supplementary Tables 2, 5 and 6**), including in representative AR-driven CRPC (**ARPC**), neuroendocrine prostate cancer (**NEPC**), neuroendocrine-like bladder cancer and ASCL1-active SCLC (r = 0.85–0.93; **Fig. 4a**). In CRPC-RA, predictions reproduced dominant androgen receptor and neuroendocrine lineage programs determined by tumor RNA-seq^1^ (**Fig. 4b**), while in patients with multiple metastases,^31^ greater RNA-seq heterogeneity correlated with higher Proteus error (ρ = 0.43, p = 0.013) and lower predictive correlation (ρ = −0.46, p = 0.009) consistent with the challenge of representing spatially heterogeneous disease (**Supplementary Fig. 8a; Supplementary Table 3**). Proteus-derived enrichment scores also correlated with RNA-seq for neuroendocrine (NE.10, r = 0.84), AR-associated (ARG.10, r = 0.81) and proliferation signatures (CCP.31, r = 0.60; **Fig. 4c; Supplementary Fig. 8b; Supplementary Table 3**).^32,33^ Applying the PAM50 classifier^34^ recapitulated Basal (r = 0.66) and Luminal A (r = 0.60) signatures, with weaker Luminal B performance (r = 0.24; **Supplementary Fig. 8c–e**).

**Figure 4.**
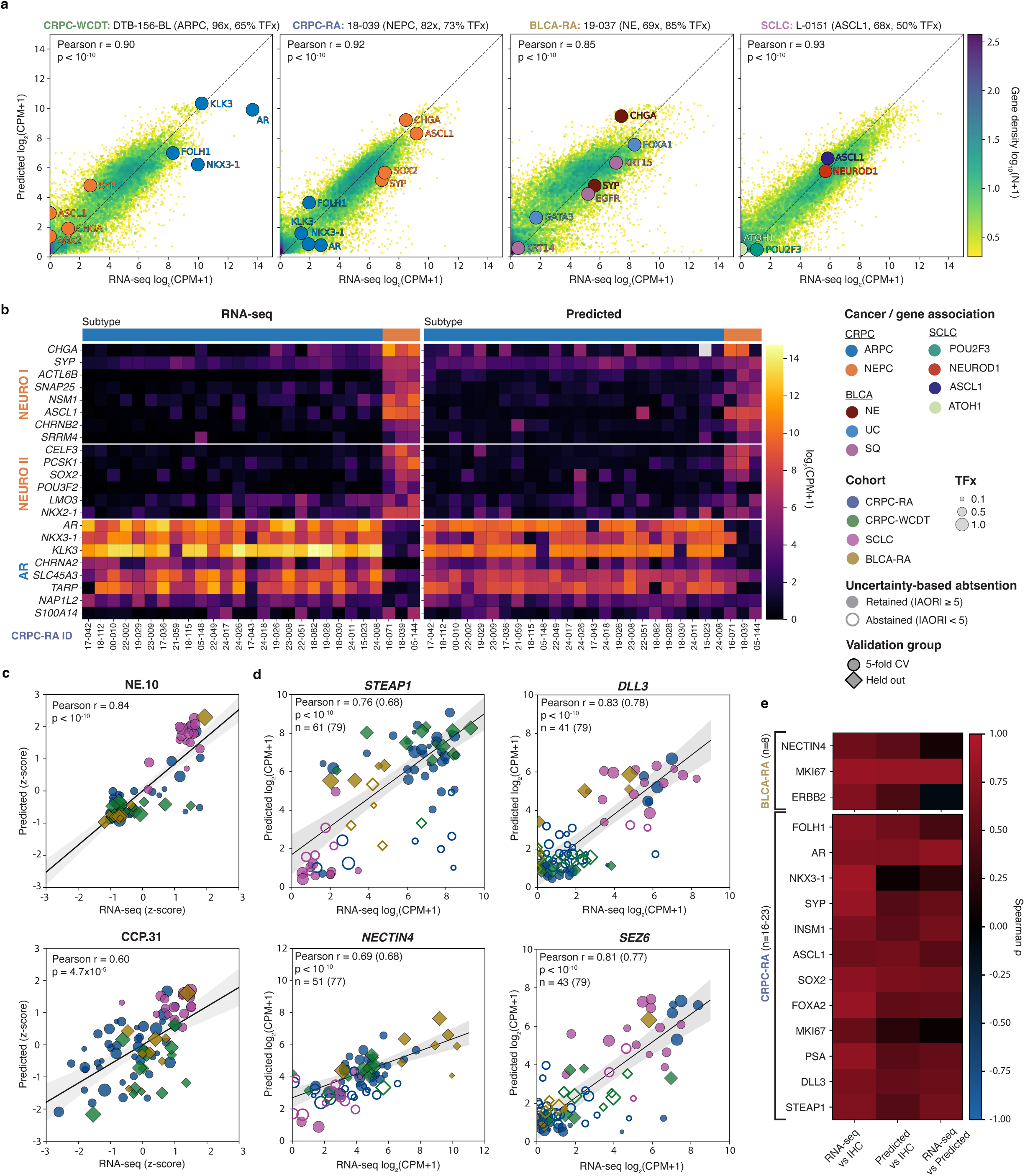
Recovery of tumor phenotype, pathway activity and therapeutic target expression from patient cfDNA. **a)** Representative transcriptome-wide comparisons between matched tumor RNA-seq and Proteus-predicted expression for individual samples: CRPC-WCDT DTB-156-BL, n=18,524 genes; CRPC-RA 18-039, n=18,528 genes; BLCA-RA 19-037, n=18,541 genes; and SCLC L-0151, n=17,848 genes. Points denote individual genes and are colored by local gene density. Selected phenotype-defining genes are labeled and colored by cancer or subtype association. **b)** Heatmaps comparing tumor RNA-seq (left) and Proteus-predicted (right) expression values for CRPC-RA patient samples with pure molecular phenotype (n = 27, **Methods**) across phenotype-defining gene sets: NEURO I, NEURO II, and AR. CRPC tumor subtypes are indicated at the top. **c)** Comparison of RNA-seq-derived and Proteus-derived ssGSEA z-scores for the NE.10 neuroendocrine signature (top) and CCP.31 cell-cycle proliferation signature (bottom) across patient cohorts (CRPC-RA, n = 44; CRPC-WCDT, n = 13; SCLC, n = 16; BLCA-RA, n = 8). Lines denote ordinary least-squares fits for visualization; shaded bands denote the 95% confidence interval around the fitted line. Reported associations use Pearson’s r. **d)** Comparison of RNA-seq and Proteus-predicted expression for STEAP1, DLL3, NECTIN4, and SEZ6 across matched patient cohorts. Lines denote ordinary least-squares fits for visualization; shaded bands denote the 95% confidence interval around the fitted line. Reported associations use Pearson’s r; total n and correlations are indicated both after and before AOR-based abstention, with pre-abstention values in parentheses. Circles indicate 5-fold CV results and diamonds held-out patient samples; hollow points indicate abstained predictions. **e)** Heatmap of Spearman correlations comparing tumor RNA-seq, Proteus-predicted expression and tumor immunohistochemistry for therapeutic and phenotype-associated markers in BLCA-RA and CRPC-RA cohorts. Tumor RNA-seq and IHC values were summarized as the mean across sampled biopsies where multiple lesions were available. Sample size varied by marker according to IHC availability; exact marker-level n values and Spearman correlation statistics are reported in **Supplementary Table 3**.

Next, we evaluated Proteus predictive accuracy for clinically relevant cell-surface targets. *STEAP1* is an emerging target for antibody–drug conjugate and cellular therapy strategies in prostate cancer.^35–37^ Predicted *STEAP1* expression correlated with tumor RNA-seq (r = 0.68), improving with AOR-based abstention (r = 0.76; **Fig. 4d**). Proteus also recovered *NECTIN4*, the target of enfortumab vedotin in bladder cancer^38,39^ (r = 0.69) and neuroendocrine tumor targets *DLL3*^40,41^ (r = 0.83) and *SEZ6*^42^ (r = 0.81; **Fig. 4d**). Comparing Proteus-predicted expression, tumor RNA-seq and tumor immunohistochemistry (**IHC**) in BLCA-RA and CRPC-RA revealed marker-dependent agreement, with Proteus-IHC associations generally following the direction of RNA-seq-IHC associations (**Fig. 4e; Supplementary Table 3; Methods**). Because Proteus infers a chromatin-defined transcriptional state, whereas RNA-seq and IHC measure transcript and protein abundance, some marker-dependent discordance is expected. Together, these findings support cfDNA-based inference for molecular phenotype characterization and therapeutic target activity for many clinically relevant genes.

### Proteus enables functional cfDNA profiling of clinical treatment states in mCRPC

We next evaluated whether Proteus predictions could support biological characterization and precision-oncology applications in clinical cohorts without matched tumor transcriptomes. First, we analyzed a lower-depth CRPC cohort classified as ARPC or NEPC by tumor histology (CRPC-UW, n = 47; depth = 15–32x; TFx = 0.03–0.90) using uncertainty-aware gene set enrichment analysis (**Methods**). Proteus recovered established lineage programs directly from cfDNA, including ARG.10 in ARPC and NE.10 and proliferative/developmental programs in NEPC cases (**Fig. 5a**). The Proteus-derived NE.10 score further distinguished pure NEPC from ARPC (AUROC = 0.90) and tumors with any neuroendocrine component from pure ARPC (AUROC = 0.77; **Supplementary Fig. 9a,b; Supplementary Table 4**).

**Figure 5.**
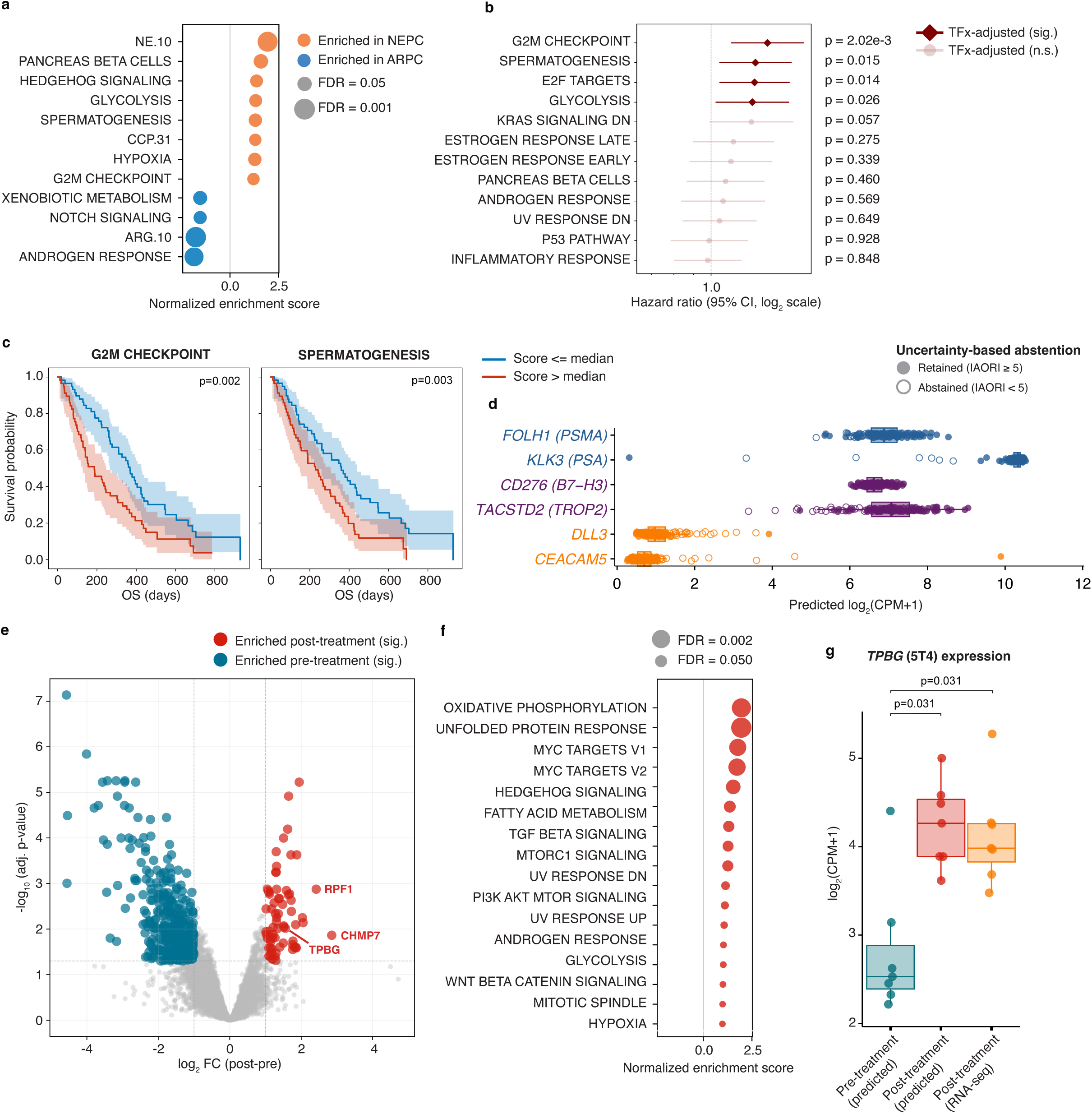
Proteus enables cfDNA-based molecular phenotyping, prognosis and longitudinal monitoring in clinical mCRPC cohorts. **a)** Gene set enrichment analysis of Proteus-predicted expression from cfDNA in clinically annotated CRPC-UW samples classified as ARPC or NEPC by tumor histology/IHC. Differential expression used limma-trend with Proteus predictive means and PI90-derived precision weights, followed by GSEA on Hallmark, ARG.10, NE.10 and CCP.31 gene sets. P values were adjusted by Benjamini–Hochberg FDR; only gene sets with FDR < 0.25 are shown. **b)** TFx-adjusted Cox proportional-hazards analysis of baseline Proteus-derived Hallmark pathway scores in patients treated with ^177^Lu–PSMA-617. Among 158 treated patients, 120 with screening TFx > 3% were selected; 115 with complete pathway-score, TFx and survival data were included in the Cox analysis, with 96 deaths. Each pathway was evaluated in a separate model containing the pathway score and TFx as covariates; both covariates were standardized before model fitting. Points show hazard ratios for overall survival per 1-s.d. increase in pathway score, and horizontal lines show 95% Wald confidence intervals. Displayed p-values are two-sided Wald tests of the pathway coefficient. Filled diamonds denote nominal p < 0.05 and are not based on multiple-testing-adjusted significance; BH-adjusted q values calculated across all 50 Hallmark pathways are reported in **Supplementary Table 4**. **c)** Kaplan–Meier curves for overall survival in patients stratified by median baseline Proteus-derived G2M checkpoint and spermatogenesis pathway scores (n = 120). Higher pathway scores were associated with shorter overall survival. P-values are from log-rank tests; shaded bands indicate 95% confidence interval. **d)** Baseline Proteus-predicted expression of selected ARPC-associated and NEPC-associated clinical or therapeutic target genes in the ^177^Lu-PSMA-617 cohort (CRPC-Pluvicto, n = 120). ARPC-associated genes include FOLH1/PSMA, KLK3/PSA, CD276/B7-H3, TACSTD2/TROP2, and NEPC-associated genes include DLL3 and CEACAM5. Each point represents one cfDNA sample/patient at baseline, with hollow points indicating abstained predictions based on uncertainty thresholding (|AOR| < 5). **e)** Paired differential expression analysis (predictive interval-weighted; **Methods**) of Proteus-predicted expression comparing post-treatment cfDNA samples with pre-treatment cycle 1 samples in patients treated with ^177^Lu-PSMA-617 (CRPC-Pluvicto, n = 7). Red points indicate genes significantly enriched post-treatment, and blue points indicate genes significantly enriched pre-treatment. Differential expression was tested using limma-trend with patient-paired design and PI90-derived precision weights; p-values were adjusted by Benjamini–Hochberg FDR. **f)** Gene set enrichment analysis of genes ranked by post-treatment versus pre-treatment differential expression from paired cfDNA samples. Post-treatment samples were enriched for metabolic, stress-response and growth-associated pathways. The top 16 post-treatment enriched gene sets are shown. **g)** Predicted expression of post-treatment-enriched candidate target gene TPBG from paired cfDNA samples before and after ^177^Lu-PSMA-617 treatment, compared with matched post-treatment tumor RNA-seq. Post-treatment cfDNA predictions were significantly increased relative to pre-treatment predictions and were concordant with post-treatment RNA-seq. P values were calculated by two-sided paired Wilcoxon signed-rank tests.

We then analyzed the prognostic utility of baseline expression profiles derived from cfDNA in 120 patients prior to receiving ^177^Lu–PSMA-617, a PSMA-targeting radioligand (CRPC-Pluvicto, depth = 21–70x, TFx = 0.03–0.81).^43,44^ As TFx is a known prognostic factor in advanced prostate cancer,^45^ TFx-adjusted Cox models were utilized for 115 patients with complete data. Higher G2M-checkpoint (HR = 1.40, 95% CI 1.13–1.74; p = 0.0020), spermatogenesis (HR = 1.30, 95% CI 1.05–1.61; p = 0.0147), E2F-target (HR = 1.30, 95% CI 1.05–1.60; p = 0.0139) and glycolysis scores (HR = 1.28, 95% CI 1.03–1.59; p = 0.0258) were nominally associated with shorter survival (**Fig. 5b**; **Supplementary Table 4**). G2M-checkpoint and spermatogenesis scores also stratified survival (**Fig. 5c**); we note that the Hallmark spermatogenesis set includes proliferation and chromatin-regulatory genes generally implicated in aggressive prostate cancer, including *AURKA*, *BUB1*, *CDK1* and *EZH2*. Proteus also predicted active expression of FOLH1/PSMA, KLK3/PSA and alternative surface targets CD276/B7-H3 and TACSTD2/TROP2, whereas NEPC-associated DLL3 and CEACAM5 were generally lower or inactive (**Fig. 5d**).^46–48^ Thus, the adverse pathway associations did not coincide with either target loss or a cohort-wide neuroendocrine transition.

Finally, seven patients with paired pre-and post-treatment cfDNA samples were used for a hypothesis-generating analysis of potential treatment-emergent states. Differential expression analysis revealed a marked post-treatment shift, and incorporating Proteus 90% PIs as precision weights increased the number of significant pre-and post-treatment-enriched genes, from 308 to 373 and 59 to 68, respectively (**Fig. 5e; Supplementary Table 4**). Post-treatment samples showed increased CHMP7 and RPF1 activity and enrichment of stress-response, metabolic and plasticity-associated programs (**Fig. 5f**). TPBG, a cell-surface protein linked to WNT signaling, epithelial–mesenchymal transition and lineage plasticity, was also increased after treatment, with the direction of change supported by matched post-treatment tumor RNA-seq (**Fig. 5g**).^49,50^ Although these paired analyses are limited by sample size, they nominate TPBG and the accompanying stress, metabolic, and plasticity programs as candidate features of treatment-emergent disease that warrant additional validation.

## DISCUSSION

Current cfDNA-based liquid biopsy assays provide a minimally invasive means of profiling tumor genotype, including the identification of genomic drivers and alterations associated with treatment response. Yet precision oncology increasingly relies on understanding tumor molecular phenotype and surface target expression that genomic profiling alone cannot capture. Herein, we have introduced two complementary tools for this purpose: Triton, a standalone method for profiling cfDNA fragmentation and inferred chromatin organization, and Proteus, a biologically informed deep-learning model that predicts individual tumor gene expression from standard-depth cfDNA WGS. Together, these methods combine local sequence context with spatial and regional coverage, nucleosome-positioning and fragmentation signals, without relying on specialized assays or targeted selection of specific regions.^11,12,51^ Model interpretation showed that Proteus learned established transcription-associated patterns, including promoter depletion near active TSSs, localized signals around the +1 nucleosome,^13,14,20^ increased fragment-length heterogeneity and altered nucleosome phasing.^16,19,21^ Gene-level and gene-variance-weighted metrics that better capture clinically relevant, patient-specific expression variation demonstrated that Proteus outperformed both a prior fragmentomics-based method^11^ and a simpler machine learning approach across admixtures, external cohorts and an unseen cancer type.

We demonstrated the utility of Proteus across molecular subtyping, therapeutic-target assessment and pathway-level analysis in clinical cohorts with and without matched tissue RNA-seq. In cohorts with RNA-seq, Proteus recovered dominant lineage programs, coordinated expression signatures and individual cell-surface targets across multiple cancer types. Predictive intervals and activity odds ratios unique to Proteus tracked error in unseen populations, and withholding low-confidence predictions further improved gene-activity calls, supporting the reporting of model-derived uncertainty alongside inferred expression. In cohorts without RNA-seq, inferred expression identified lineage-associated pathways and nominated transcriptional programs and therapeutic targets associated with treatment-emergent disease. These vignettes illustrate how cfDNA-derived expression profiles could enable longitudinal assessment of tumor state, treatment response, resistance and plasticity beyond measurements of tumor fraction or genomic alterations alone.

Proteus establishes a deep-learning framework for inferring tumor gene expression from standard cfDNA WGS. However, its performance remains constrained by training-data diversity, particularly for poorly represented genes, low-depth or low-TFx samples and heterogeneous tumor mixtures. Its confidence estimates do not remove these limitations, but provide a means to identify and account for them. Moreover, Proteus infers chromatin-associated transcriptional states and cannot yet resolve individual lesions, subclones, or immune and tissue contributions within mixed cfDNA. Larger and more diverse training datasets should broaden the set of reliably predicted genes and improve generalizability, while incorporating methylation, histone modifications and other cfDNA modalities could support tissue-, immune-and lesion-level deconvolution. Together, Triton and Proteus constitute a foundation for functional liquid biopsies, moving cfDNA analysis beyond tumor detection or mutational profiling and towards minimally invasive, longitudinal measurements of tumor regulatory states and the identification of therapeutic targets that reflect tumor phenotype.

## METHODS

### PDX mouse models

#### LuCaP CRPC PDX Models

Twenty-five LuCaP PDX models with line-matched cfDNA sequencing and tumor tissue RNA-seq were previously described^16,52^. PDXs were propagated *in vivo* in male NOD/SCID IL2R-gamma-null mice (NSG; cat. #005557), with tumor collection for initial establishment approved by the University of Washington (UW) Human Subjects Division Institutional Review Board (IRB #2341). Phenotypes were determined through histopathology by at least two expert pathologists and validated using transcriptome-derived marker expression scores^33,53,54^.

Blood was collected via cardiac puncture from mice bearing tumors of 300–1,400 mm³. Samples were collected into Sarstedt K3-EDTA microtubes and processed within 4 h, undergoing sequential centrifugation at 2,500 × g for 10 min and 16,000 × g for 10 min (room temperature). Plasma from 4–8 mice was pooled per PDX line, transferred to screw-cap cryovials, and stored at −80 °C for subsequent cfDNA isolation. All animal procedures were conducted in accordance with NIH guidelines and approved by the Fred Hutchinson Cancer Center IACUC (protocol #1618).

Sequencing libraries were built from 50 ng of input cfDNA with the KAPA HyperPrep kit, using nine PCR cycles, SPRI-bead cleanup, and KAPA UDI dual-indexed adapters. Libraries were quantified, equalized, pooled for multiplexing, and sequenced on an Illumina HiSeq 2500 (200-cycle kit) at the Fred Hutchinson Genomics Shared Resources or on an Illumina NovaSeq employing S4 flow cells (300-cycle kit) through the Broad Institute Genomics Platform Walkup-Seq Services. To keep read lengths consistent, the NovaSeq output was trimmed to 200 cycles, producing 100 bp paired-end FASTQ files that match the HiSeq data.

Published bulk tumor RNA-seq data from the LuCaP PDX tumors were processed as described previously^55^. Reads were aligned to the hg38 human genome using STAR v2.7.3a. PDX data were also aligned to the mm10 mouse genome. PDX sequencing reads deriving from mouse were removed using XenofilteR,^56^ and gene-level abundance was quantitated using GenomicAlignments.

#### SCLC PDX Models

Twenty-five SCLC and non-SCLC PDX models with line-matched cfDNA sequencing and tumor tissue RNA-seq were previously described^10^. Models were established from circulating tumor cells or tumor tissue^57–60^ and propagated *in vivo* in male NOD/SCID IL2R-gamma-null (NSG) mice (cat. #005557). Blood was collected by cardiac puncture from animals bearing flank tumors; blood from up to five mice of the same model was pooled when available. Samples were processed within 4 h of collection, undergoing centrifugation at 2,500 × g for 10 min at room temperature to separate plasma, followed by a second centrifugation of plasma at 13,200 × g for 10 min to remove residual debris. Plasma and, when collected, buffy coat fractions were stored at −80 °C for subsequent DNA isolation. All animal procedures were performed in accordance with institutional guidelines and approved protocols.

Genomic DNA was extracted from buffy coat using the QIAamp DNA Blood Mini Kit (QIAGEN, 51104), while cfDNA was isolated from 0.5–4 ml of plasma using either the QIAamp MinElute ccfDNA Mini Kit (QIAGEN, 55204) or QIAamp Circulating Nucleic Acid Kit (QIAGEN, 55114), following manufacturer instructions. DNA concentration was measured using the Qubit dsDNA HS Assay Kit (Thermo Fisher Scientific, Q32851), and size distribution assessed on a Bioanalyzer using the DNA 1000 or HS DNA 1000 Kit (Agilent, 5067-1504 or 5067-4626). Plasma cfDNA libraries were prepared with either the NEBNext Ultra II DNA Library Prep Kit for Illumina (New England Biolabs, E7645 and E7335/E6609) or the xGen cfDNA & FFPE DNA Library Prep MC Kit with unique dual index primers (Integrated DNA Technologies, 10006203 and 10005922). For buffy coat genomic DNA, samples were mechanically sheared to ∼250 bp modal size using a Bioruptor (Diagenode; low power, 30 s on / 90 s off, 4 °C, 30 min), followed by library construction using the xGen cfDNA & FFPE DNA Library Prep MC Kit.

RNA was extracted from PDX tumor tissue or cell pellets using TRIzol reagent, and RNA-seq libraries were prepared from 500 ng total RNA with the NEBNext Ultra RNA Library Prep Kit for Illumina (New England Biolabs, E7530L). Sequencing was performed on an Illumina HiSeq 2500 instrument with 50 bp single-end reads. Reads were aligned using STAR v2.7.1a in two-pass mode to both the human genome (hg38, GENCODE v31) and mouse genome (mm10, GENCODE vM23), followed by removal of mouse-derived reads with XenofilteR. QC metrics were generated with FastQC v0.11.9 and RSeQC v4.0.0. Gene-level counts were obtained in an unstranded manner using FeatureCounts (Subread v1.6.5).

### cfDNA sequence alignment and mouse subtraction

All cfDNA reads were re-aligned to the Broad GRCh38/hg38 human reference genome with BWA-MEM v0.7.17^61^; the Snakemake workflow and configuration are available at https://github.com/GavinHaLab/fastq_to_bam_paired_snakemake. Alignment and coverage metrics were computed with Picard CollectAlignmentSummaryMetrics and CollectWgsMetrics (GATK/Picard v4.1.8.1). For PDX ctDNA WGS datasets, mouse-derived reads were removed as previously described^10,16,62^.

### Human cohorts

#### HD (Healthy Donor) cohorts

The HD-L cohort comprised ten donors with matched plasma cfDNA and PBMC RNA-seq. Two 10 ml EDTA blood tubes per donor were obtained from Bloodworks Bio (Seattle, WA; 4500-01), transported on ice and processed within 2 h. Blood was centrifuged at 500 × g for 15 min at 4 °C, and plasma was transferred and centrifuged at 15,000 × g for 10 min at 4 °C. Plasma aliquots were stored at −80 °C. cfDNA was extracted from up to 4 ml plasma with the QIAamp Circulating Nucleic Acid Kit (QIAGEN, 55114). PBMC pellets were lysed in RLT buffer containing β-mercaptoethanol and stored at −80 °C; RNA was extracted with the QIAGEN RNeasy Kit with on-column DNase digestion (79254).

The HD-U cohort comprised plasma from ten donors without matched RNA-seq (Research Blood Components, Watertown, MA; 1-009). Plasma was separated at 1,600 × g for 10 min at room temperature, shipped frozen and stored at −80 °C. After thawing, samples were centrifuged at 15,000 × g for 10 min. cfDNA was extracted from 4 ml plasma using the QIAamp Circulating Nucleic Acid Kit and eluted in 55 µl Buffer AVE. Concentration was measured using the Qubit 1X dsDNA High Sensitivity Assay (Thermo Fisher Scientific, Q33230), and fragment sizes were assessed using the Agilent Cell-free DNA ScreenTape assay on a 4200 TapeStation.

### CRPC-RA and BLCA-RA cohorts

Participants in the CRPC-RA and BLCA-RA rapid-autopsy cohorts provided written informed consent under the UW Genitourinary Cancer Biorepository protocol (IRB 2341), including consent for rapid autopsy. Tissues were procured within 15 h of death for CRPC-RA (median, 5 h) and within 11.5 h for BLCA-RA (median, 3.4 h). Cardiac or peripheral blood was collected at the start of autopsy into serum or EDTA tubes, centrifuged at 1,600 × g for 15 min, aliquoted and stored at −80 °C. cfDNA was extracted from 0.5-2 ml plasma or serum as described for HD-U. For CRPC-RA, 44 patients contributed cfDNA and a total of 104 metastatic, flash-frozen tumors (one to five per patient) for RNA-seq. RNA was isolated from OCT-embedded metastases using RNA STAT-60 after tumor-enriched 1-mm core sampling or cryostat sectioning guided by hematoxylin and eosin staining. Seven patients also belonged to the CRPC-Pluvicto cohort and contributed matched post-treatment rapid-autopsy cfDNA and tumor RNA-seq; their post-treatment expression estimates were generated using patient-held-out cross-validation. Individual metastases were assigned ARPC or NEPC phenotype based on either IHC or RNA-seq centroid-based classification where available, as previously described;^1^ 27 patients were defined as having pure ARPC or NEPC based on all queried metastases having the same phenotype classification.

For BLCA-RA, eight patients contributed cfDNA and tumor specimens, including seven primary tumors (four formalin-fixed paraffin-embedded (FFPE) and three flash-frozen) and 28 flash-frozen metastases (two to five per patient). Primary and metastatic tissues were embedded in OCT or processed as FFPE material. Histology was assigned by expert pathology review.^63,64^

### SCLC, CRPC-WCDT, and CRPC-UW cohorts

The patient SCLC cohort was described previously.^10^ Blood was collected into K2EDTA tubes and centrifuged at 2,500 × g for 10 min at room temperature. Plasma was transferred, centrifuged at 13,200 × g for 10 min and stored at −80 °C. Buffy coat, when collected, was also stored at −80 °C. Genomic DNA and cfDNA were isolated and assessed using the same procedures described for the SCLC PDX cohort.

CRPC West Coast Dream Team (CRPC-WCDT) samples with matched cfDNA and tumor RNA-seq were described previously^26^ and were obtained with permission directly from the study authors.

The CRPC-UW WGS cohort was described previously.^16^ Briefly, 31 men with metastatic castration-resistant prostate cancer (mCRPC) enrolled under UW IRB CC6932 between 2014 and 2021 and provided 61 plasma samples with written informed consent. Following ultra-low-pass WGS screening, 47 samples from 27 patients with ichorCNA-estimated tumor fraction greater than 3% underwent high-depth WGS. Only high-depth samples and associated clinical histology were analyzed here.

### CRPC cohort receiving ^177^Lutetium-PSMA-617 (CRPC-Pluvicto)

Patients receiving ^177^Lutetium-PSMA-617 (Pluvicto) provided written informed consent under UW IRB CC6932 (NCT01050504). The source cohort comprised 158 patients with metastatic castration-resistant prostate cancer who met United States FDA-label clinical criteria for treatment with ¹⁷⁷Lu-PSMA and had pretreatment PSMA-PET/CT imaging and a baseline blood sample available for cfDNA analysis. Of these, 120 with ichorCNA-estimated TFx > 3% on preliminary ultra-low-pass WGS underwent deeper WGS and were included here; ultra-low-pass data were used only for sample selection. Patients were treated with up to six cycles of ¹⁷⁷Lu-PSMA, each 7,400 MBq (200 mCi) ± 10%. During each cycle, patients were clinically evaluated by a group of oncologists with expertise in prostate cancer management. Peripheral blood samples for cfDNA analysis were collected immediately before cycle 1 (C1) into Streck Cell-Free DNA BCT or K2EDTA tubes and processed within 36 h or 4 h, respectively. Blood was centrifuged at 2,500 × g for 10 min, and plasma was transferred and centrifuged at 3,000 × g for 15 min. Plasma and buffy-coat fractions were aliquoted and stored at −80 °C. cfDNA was purified from 2 ml plasma by Broad Clinical Laboratories using the QIAsymphony DSP Circulating DNA Kit (QIAGEN, 937556), eluted in TE buffer and quantified with PicoGreen. Overall survival (OS) was defined from first ¹⁷⁷Lu-PSMA treatment to death from any cause, with patients censored at last known follow-up through the data cutoff of August 15, 2025.

### cfDNA library preparation and sequencing

For internal HD-U, BLCA-RA, CRPC-RA and CRPC-Pluvicto samples, library preparation and sequencing were performed by Broad Clinical Laboratories. cfDNA input (25–52 ng in 50 µl TE) was prepared with the KAPA HyperPrep Kit (KK8504) and IDT xGen UDI-UMI duplex adapters without prior shearing. Libraries were purified with AMPure XP beads, quantified using PicoGreen and normalized. Ultra-low-pass libraries were initially generated and sequenced as paired 151-bp reads on NovaSeq 6000 SP flow cells. Libraries selected for deeper sequencing used in this study were repooled after qPCR normalization and sequenced as paired 151-bp reads on NovaSeq X 25B flow cells.

For HD-L, libraries were prepared from 100 ng cfDNA using the IDT xGen cfDNA & FFPE DNA Library Prep v2 MC Kit and xGen indexing primers. Libraries were assessed by Qubit and TapeStation, pooled at equal molarity and sequenced as paired 150-bp reads on a NovaSeq X Plus.

### ctDNA tumor–normal admixtures

In silico admixtures for model training and benchmarking (PHT and PHS sets) were generated using all available PDX (LuCaP CRPC n = 25, SCLC = 25) and HD (HD-L (with RNA-seq) n = 10, HD-U (no RNA-seq) n = 10) samples. All mixtures comprised a single PDX and single HD sample. Training admixtures (PHT; n = 80) combined the 40 development PDX models with HD-L donors and their matched PBMC RNA-seq. Benchmarking admixtures (PHS) combined ten held-out PDX models (five CRPC and five SCLC) with HD-U donors; neither those PDX models nor any HD-U donor contributed to PHT or model development. For the Figure 2b benchmark, PHS input samples were downsampled before mixing to generate 120 admixtures at 5× and 10× depth and tumor fractions of 1%, 3%, 5%, 10%, 30% and 50% (n = 10 per depth–TFx condition). Complete mixture descriptions are available in Supplementary Table 1. BAM files were merged with SAMtools; the full Snakemake workflow for merging and down-sampling is available at https://github.com/GavinHaLab/Admixtures_snakemake.

### RNA-seq processing

Published data of bulk tumor RNA-seq for LuCaP PDX models, SCLC PDX models, the SCLC patient cohort, and the CRPC-WCDT cohort were previously described^10,26,55^. Bulk tumor RNA-seq for the CRPC-RA cohort was sequenced and aligned as described previously^55^. In brief, sequencing reads were mapped to the hg38 human genome using STAR v2.7.3a. All subsequent analyses were performed in *R*, with gene-level abundance quantitated using *GenomicAlignments*.

For the HD-L cohort, RNA was quantified using a Lunatic spectrophotometer (Unchained Labs, Pleasanton, CA) and RNA quality was confirmed using an Agilent 4200 TapeStation (Agilent Technologies, Santa Clara, CA). Sequencing libraries were prepared using the TruSeq Stranded mRNA kit (Illumina, Inc., San Diego, CA) using 250 ng total RNA input. Library fragment size distributions were confirmed using Agilent 4200 TapeStation and library yields were determined using Life Technologies’ Invitrogen Qubit 2.0 Fluorometer (Thermo Fisher, Waltham, MA). Sequencing libraries were pooled in equimolar ratios and sequenced on an Illumina NovaSeq X Plus (Illumina, Inc, San Diego, CA) over one lane of a 10B-300 flow cell employing a paired-end, 50-base read length sequencing configuration.

For the BLCA-RA cohort, RNA was isolated from 42 flash-frozen autopsy specimens embedded in OCT using a combined protocol of RNA STAT-60 (Tel-Test, Friendswood, TX) and the RNeasy Mini Kit with in-solution DNase digestion prior to column purification (Qiagen, Germantown, MD). For eight FFPE surgical specimens, RNA was extracted with the AllPrep DNA/RNA FFPE Kit (Qiagen, Germantown, MD). RNA quality was assessed by RNA integrity number (RIN) on an Agilent Bioanalyzer (Agilent Technologies, Santa Clara, CA). For RNA sequencing, libraries were prepared from 300 ng total RNA using the TruSeq RNA Exome Sample Preparation Kit following the manufacturer’s instructions (Illumina, San Diego, CA). Barcoded libraries were pooled and sequenced on an Illumina HiSeq 2500 to generate 50 bp paired-end reads.

Both internal and external RNA-seq cohort gene-level counts were mapped or re-mapped to the MANE Select^65^ v1.3 primary transcripts based on either ENTREZ IDs or gene symbols, as available. Sample-level raw counts for all cohorts underwent Trimmed Mean of M-values (TMM) re-scaling as implemented in *edgeR*^66^ to normalize differences in library sizes. Normalized raw counts were then converted into Counts Per Million (CPM) and subsequently into log_2_(CPM+1) for use as training labels and in all downstream RNA-seq analyses. For samples with RNA-seq from multiple tissue biopsies, log_2_(CPM+1) values were averaged across biopsies to obtain a single representative transcriptome per sample. CPM units were chosen for training as they are compatible with standard RNA-seq workflows and can be converted to length-normalized TPM values when transcript lengths are available. RNA-seq expression values were deliberately not batch-corrected using *ComBat* or related cross-cohort empirical Bayes procedures to guard against data leakage in model development and benchmarking.

### Patient tumor fraction estimation

For external data sets (SCLC, CRPC-UW and CRPC-WCDT cohorts) the provided tumor fraction estimates were used. Elsewhere, tumor fraction was estimated for cfDNA WGS samples using the *TITAN*^67^ pipeline (v1.15.0; https://github.com/GavinHaLab/TitanCNA_SV_WGS, commit ID: bedbd76). For four samples in the BLCA-RA cohort and all samples in the CRPC-RA cohort with matched tissue normal WGS, standard tumor–normal paired analysis was performed. For the remaining four BLCA-RA samples and cohorts without matched WGS normals, a tumor-only pipeline was applied. *TITAN*-derived results were manually curated to determine the optimal solution for each sample, from which tumor fraction estimates were obtained. In the CRPC-Pluvicto cohort, tumor content in patient plasma cfDNA was quantified with *ichorCNA*^68^ using default settings and 1 Mb bins. ichorCNA’s default tumor-fraction output was manually reviewed, and optimal solutions were selected based on expert curation.

### Immunohistochemistry (IHC)

Tumor tissues from bladder and prostate rapid autopsy cases (CRPC-RA and BLCA-RA cohorts) were previously evaluated for target protein expression using validated immunohistochemical assays. When multiple metastasis IHC values were available, the mean was used. Details regarding the sample cohorts, immunohistochemistry protocols, and expert pathologist assessment have been previously described^31,48,64,69^.

### Triton nucleosome profiling and fragmentation pattern analysis

#### Overview

*Triton* converts mapped cfDNA reads into nucleotide-resolution “spatial” signal profiles and matched region-level biomarkers. The user supplies a BED-style coordinate set as either (i) single genomic regions, for example full-length gene bodies (“region” mode), (ii) fixed-width windows centered on an index such as transcription-start sites (“window” mode, default ±1 kb), or (iii) collections of windows that are first “piled-up” across many instances of the same regulatory element (“composite-window” mode). For every site/region or composite, *Triton* extracts properly-paired fragments (default 15–500 bp) from the BAM/CRAM, applies per-fragment corrections (see below), and finally reports both bp-resolution tracks and region-level fragmentation and nucleosome-phasing metrics, summarized into a tab-separated feature table. In this study, MANE Select v1.3 transcription-start-site (TSS) windows (TSS ±1,000 bp; window mode) and full transcript bodies (region mode) were analyzed. Gene-body intervals spanned the genomic extent of each MANE Select transcript, including exons and introns; the downstream portion of the TSS window therefore overlaps the 5′ gene body when transcript length permits. Exon–intron junctions were not modeled separately.

For each region, *Triton* produces 12 spatial channels: GC-corrected fragment depth, phased-nucleosome (PN) signal, fragment-end density, fragment-end orientation asymmetry, subnucleosomal-fragment ratio, fragment-length Shannon entropy, fragment-length Gini-Simpson diversity, called PN peaks, and four reference-nucleotide channels. Eleven scalar summaries are calculated independently for promoter and gene body: fragment-length mean, standard deviation, skewness and kurtosis; subnucleosomal ratio; Shannon entropy; Gini-Simpson diversity; PN amplitude, mean spacing and periodicity score; and mean depth (**Supplementary Fig. 1a**).

#### Fragment filtering and GC correction

After removing PCR duplicates, reads with mapping quality < 20, and fragments with length outside 15–500 bp, GC correction is performed. *Triton* adopts the fragment-level bias tables produced by *Griffin*^17^ (fragment length × GC%) but will accept any bias-correction file in the same format. During iteration over every fragment within *Triton* processing of an individual site, the bias factor is looked up (fragment_bias) and the fragment is accepted only if 0.05 (0.10 for region mode) < bias < 10. Its contribution to depth is then rescaled by 1 / fragment_bias. This bias correction is not applied when calculating fragment length distributions or end coverage, wherein the 5’ and 3’ ends of all fragments passing filtering are piled up.

#### Fragment-length re-weighting for nucleosome-center localization

To produce the phased-nucleosome (PN) signal and related features, fragments ≥146 bp are re-weighted to reflect the empirical probability that a given base along each fragment coincides with a histone dyad. This is achieved with a pre-computed weight matrix (NCDict.pkl), where each row encodes the displacement density for one fragment length, meaning longer fragments contribute less sharply than core-sized fragments and reflect underlying uncertainty in wrapping symmetry. The weight matrix was formed by running “nc_dist.py” on healthy donor cfDNA samples, a simplified version of *Triton* which runs in “composite-window” mode and returns arrays with counts of displacements between fragment centers and the central site. A set of 186 sites representing high-confidence nucleosome centers was used; iNPS^70^ peak BED files across multiple tissues were downloaded from NucMap^70,71^ (https://download.cncb.ac.cn/nucmap/organisms/latest/Homo_sapiens/data_type/iNPS_peaks/, retrieved all BED files as of 2023-07-20) and sites with “MainPeak” identity intersected using BEDTools v2.30.0. Sites with >50% overlap across all experiments were kept, resulting in 186 sites with nucleosome peak locations consistent across human tissues and conditions. Raw displacement profiles generated by Triton (nc_dist.py) for those sites and samples for each possible fragment length then underwent Gaussian smoothing. Finally, displacement profiles were fit to a central Gaussian and/or symmetric flanking Gaussian profile using scipy.optimize curve_fit (v1.8.0) to create the final, smoothed fragment weight matrix (visualized in **Supplementary Fig. 1b**). At run time Triton retrieves the smoothed profile from the weight matrix for each given fragment length and uses that value (divided by the fragment’s GC bias, if provided) to create the nucleosome-center signal (nc_signal) used downstream in PN signal generation.

#### Fourier filtering to obtain the phased-nucleosome (PN) signal

To generate the PN signal, the nc_signal derived from GC-corrected and fragment-length re-weighted fragments is then processed with a low-pass filter by performing a fast Fourier transform (FFT; scipy.fft v1.8.0) on the signal plus 5000 bp flanking regions used as padding. This process removes high-frequency components corresponding to periods shorter than 146 bp before the signal is reconstructed (via inverse FFT). This cutoff ensures that the signal used for downstream analyses reflects the minimum possible internucleosomal spacing, and excludes peak pileups unrelated to dominant nucleosome phasing.

#### PN-signal spatial profiles and region-level features

In addition to reporting the bp-resolution GC-corrected coverage profile and PN signal, *Triton* also performs peak calling of maxima and minima on the PN signal to localize peaks and troughs in the nucleosome signal (encoded as +1 and -1). From the PN signal and reported peaks the following region-level features are derived: *PN mean amplitude* which measures signal-to-noise ratio in nucleosome positioning (mean PN signal amplitude, based on called peaks, normalized to the mean PN signal), *PN periodicity ratio* which measures strength of phasing in two linker regimes (the ratio of mean amplitudes in the 180-210 bp periodicity to 150-180 bp periodicity ranges, based on FFT spectra), and *PN mean spacing* (mean internucleosomal distance, based on called peaks).

#### Fragmentation-based spatial profiles and region-level features

For each site, *Triton* records a histogram of observed fragment lengths (default 15–500 bp) which overlap the region at any point (“region-level”) as well as histograms for every point in the region, which include all fragments overlapping each single bp locus (“spatial”). Calculation of fragment length-derived features is performed identically based on both region-level and spatial histograms and includes: fragment length mean and standard deviation; fragment length skew and kurtosis; subnucleosomal ratio, the ratio of short/subnucleosomal (≤147 bp) to long (>147 bp) fragments; Shannon entropy (in bits); and the Gini-Simpson diversity index.

### Proteus deep learning method for gene expression prediction

#### Overview

*Proteus* is a multi-input deep neural network that predicts single gene expression from cfDNA nucleosome positioning and fragmentation pattern profiles produced by *Triton*. For all samples used in both training and prediction, *Triton* was first run on MANE Select^65^ v1.3 full transcripts (gene bodies, using region mode) and TSS ±1,000 bp windows (promoter regions, using window mode) to generate training and testing data. *Proteus* encodes each gene’s input features into two parallel probabilistic latent spaces, one intended to capture tumor-originating information, and one intended to capture background/whole-blood information, using dual variational autoencoders. A contrastive loss term encourages separation between these latents, while a learned, tumor fraction-conditioned recombination of the latents reconstructs the original scalar features, enforcing an internal self-consistency constraint. Expression is then predicted from each latent. *Proteus* is implemented in PyTorch as BioDVN, a biologically informed disentangling variational network that predicts gene-level tumor expression from cfDNA WGS.

#### Inputs and pre-processing

*Triton* outputs were compiled into per-sample Zarr stores containing TSS signals (2,000 bp × 9 channels), TSS reference sequence (2,000 bp × 4 one-hot channels), TSS scalar features (11), gene-body scalar features (11), reference scalars (3), tumor-fraction labels and tumor and background expression labels where available. The nine TSS channels comprise eight biological channels and a binary missingness mask. Missing or infinite spatial values were set to zero after constructing the mask. Depth, fragment-end density and PN signal were divided by their gene-level means and clipped to [-10, 10].

Mean region depth was transformed with log(1 + x) without standardization. Fragment-length kurtosis, PN spacing and PN amplitude were log transformed and then standardized across genes within each sample; remaining scalar features were standardized without log transformation. Residual missing scalar values were replaced by the within-sample feature mean. Genes with more than 50% missing required region-level features were excluded. Reference scalars (number of exons, mRNA size and transcript length) were log transformed and standardized across the genome. Targets were TMM-normalized log_2_(CPM + 1) values as described above; missing labels were masked in loss calculations.

#### Architecture

The full model processes each of nine TSS signal channels using grouped one-dimensional convolutions and the four nucleotide channels using a two-dimensional convolution spanning the nucleotide axis. Production convolutions used 16 filters and a 3-bp kernel. Outputs were concatenated and passed through three residual cross-channel convolutional reduction stages (interaction kernel 5, dilation 3), each followed by average pooling. The complete reduced promoter map was then flattened through a dense funnel to a 64-dimensional representation.

This representation was concatenated with the 11 TSS, 11 gene-body and three reference features. Parallel variational encoders (initial width 256; latent dimension 64; dropout 0.25) then produced Gaussian tumor and background latent distributions; reparameterized samples were used during training and latent means during inference.

An auxiliary network predicted tumor fraction (when supplied as an auxiliary label) from both latent means. Predicted tumor fraction and the latent means and log variances informed softmax weights that combined the Gaussian latents using the law of total variance. A decoder reconstructed the fused promoter and scalar representation. Ground-truth tumor fraction supervised the auxiliary predictor during training but was never supplied to the prediction path at inference.

Separate mixture-density heads mapped tumor and background latents to three-component distributions over log_2_(CPM + 1): a point mass at zero, a Gamma distribution with fixed shape *k* = 0.75 and learned rate *β*, and a Normal distribution with learned mean and standard deviation. For corresponding weights *π*_0_, *π*_G_ and *π*_N_, the point estimates were calculated as: 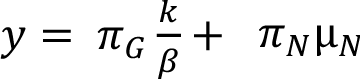 (**Supplementary Fig. 1c**).

The 5th and 95th percentiles were obtained by bisection of the mixture cumulative distribution function. The full, no-sequence and sequence-only models contained 4,512,344, 4,505,960 and 4,441,759 trainable parameters, respectively. The no-sequence ablation omitted nucleotide channels, while the sequence-only ablation used only nucleotide channels, omitted all scalar covariates, and disabled tumor-fraction supervision, latent reconstruction and sample-dependent structure losses. Ablation models used the same development samples, fixed folds, source groups, architecture widths and optimization schedule as full BioDVN; only the stated input paths and sequence-only objective differed (**Supplementary Fig. 2b**).

#### Training objective

The objective comprising both internal and external components was ℒ = ℒ_external_ + ℒ_internal_. For a predicted mixture *F* = ∑*_i_ π_i_ F_i_* and observation *y*, the analytical Continuous Ranked Probability Score (CRPS)^72^ was used:

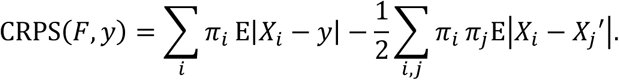

Closed forms were used where available, with the Gamma-Normal interaction using 32-node Gauss-Laguerre quadrature. CRPS values were weighted by each gene’s expression standard deviation divided by the median positive standard deviation, clipped to [0.5, 2.0] and renormalized within each batch. Admixture samples were excluded when estimating variances because their labels duplicate their parent PDX or healthy-donor labels.

For full and no-sequence models,

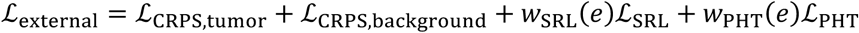

where the standardized-residual loss (SRL) was a Huber loss (*δ* = 1) after centering and scaling predictions and observations by each gene’s training-set mean and standard deviation. PHT consistency minimized, per gene, variation in tumor predictions among admixtures sharing a parent PDX and variation in background predictions among admixtures sharing a healthy donor. SRL and PHT weights were zero for five epochs and cosine-ramped over 85 epochs to 1.5 and 0.5. Optional dynamic-pair and PHT uncertainty terms had weight zero in final experiments.

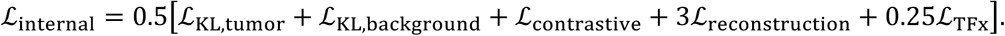

Kullback-Leibler divergence (KL) terms regularized both approximate posteriors toward a unit Normal prior. The contrastive term was max(0, *m* − *d*), where *m* = 1 and *d* was mean squared distance between sampled tumor and background latents. Reconstruction and TFx supervision used mean-squared error. The sequence-only model was trained using CRPS alone.

#### Sampling, optimization and model selection

Development samples were assigned to five fixed folds (“TDS” folds as described in **Supplementary Table 1**). Four source groups were balanced during training and validation: CRPC-RA, SCLC, prostate admixtures/PDX and SCLC admixtures/PDX, where training admixtures were generated using development PDX models and HD-L donors. The multi-source loader formed gene-conditioned 64-example batches containing two genes and 32 samples per gene, with balanced sampling across source groups. Training used 4,000 stochastic batches per epoch and seed 23; validation exhaustively evaluated held-out pairs.

Architecture, optimization and loss settings were selected by coordinated fold-parallel Optuna tuning. Final runs used a fixed 100 epoch schedule of fused AdamW, peak learning rate 1.5 x 10^-4^, no weight decay, mixed precision and gradient-norm clipping at 10. Learning rate increased linearly from 1 x 10^-5^ during the first 5.6% of scheduled steps and followed cosine decay to 1.5 x 10^-6^ over 90% of training. An exponential moving average (EMA; decay 0.995) was updated after each optimizer step; validation and checkpoints used EMA weights.

In five-fold cross-validation, four folds were used for training and one for validation. All gene– sample instances from a biological sample were assigned to the same partition; admixtures derived from the same PDX and healthy-donor parents were also grouped across partitions, where applicable. Validation occurred every five epochs and at the terminal epoch. Checkpoint selection minimized a source-level soft-maximum objective. The production model was then trained for 100 epochs on all development samples and the terminal EMA state was saved. CRPC-RA and SCLC were analyzed using patient-held-out, out-of-fold predictions; independent and discovery samples used the full-training checkpoint. For the seven CRPC-Pluvicto patients overlapping CRPC-RA, post-treatment predictions were out-of-fold, whereas pretreatment samples were absent from training but their patients contributed post-treatment samples to the full-training model. All analyses other than ablation benchmarks used the full BioDVN architecture. Training and inference were tested on NVIDIA RTX 2080 Ti and L40S GPUs; inference of approximately 18,000 genes required less than 1 min per sample. Single-GPU training of the full production configuration completed in approximately 6 h in the study environment.

#### Confidence reporting

Proteus reports predictive means, variances, mixture parameters and 90% predictive intervals. Activity confidence (activity odds ratio; AOR) was calculated from output distribution parameters as:

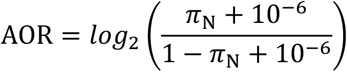

where π_N_ is the normal (active) component mixture weight. For binary gene-activity analyses, predictions with |AOR| < 5 were considered ambiguous and excluded from calls. The illustrative threshold |AOR| = 5 was selected from cross-validation error–coverage behavior before evaluation of the held-out cohorts and was not optimized on CRPC-WCDT or BLCA-RA (Supplementary Fig. 3c). Gene-specific AOR thresholds should be established based on user needs for clinical use cases. Point ssGSEA, limma and PAM50 analyses used predictive means. Predictive intervals were used as precision weights for downstream limma-based analyses (Fig. 5e; Supplementary Table 4).

#### Model interpretation

Model attribution used SHAP GradientExplainer expected gradients.^29^ The analysis sampled up to 4,096 background and 1,536 test gene-sample examples; the explainer summarized the background with at most 128 examples. Attributions were calculated for biological TSS signals, nucleotide sequence, TSS features, gene-body features and reference scalars.

For signal channel *c* and position *p*, signed polarity was calculated as:

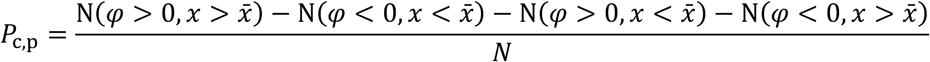

where *φ* is the SHAP value. Polarity-impact was *P*_c,p_ times median absolute SHAP value. UMAP was applied to deterministic tumor-latent means to visualize relationships with RNA-seq expression, tumor fraction, cohort, error and uncertainty (**Supplementary Figs. 5–7**).

### Benchmarking and comparator methods

To illustrate the limitation of sample-level correlation, we constructed two RNA-seq-only benchmark controls using CRPC PDX RNA-seq profiles. In the naïve control, each predicted gene value was sampled from the measured expression of the same gene in another sample, preserving gene-wise abundance distributions while destroying sample-specific biological variation. In the nearest-neighbor control, PCA was applied to RNA-seq profiles and each sample was assigned the complete expression profile of its nearest transcriptomic neighbor, excluding itself (pairings available in **Supplementary Table 2**). Both controls were evaluated using the same finite-gene masks and correlation metrics as model predictions.

The XGBoost comparator used the same Zarr stores, tumor-expression labels and fixed folds as Proteus. One pooled histogram-based regressor was fit across gene-sample rows using 25 scalar inputs (11 TSS features, 11 gene-body features and three reference features); spatial TSS signals and nucleotide sequence were excluded, and rows with non-finite tumor-expression labels were removed. Training rows were weighted by gene-expression variance calculated in the applicable training samples and clipped to [0.1, 10]. Hyperparameters were selected in 86 Optuna trials by five-fold cross-validation, with 30-round early stopping and a source-weighted gene-variance-weighted MAE objective across CRPC-RA, SCLC, prostate admixtures/PDX and SCLC admixtures/PDX. The selected absolute-error regressor used maximum depth 13, learning rate 0.0530007, minimum child weight 21.7529, row subsampling 0.8651, column subsampling 0.7642, L1 penalty 4.05 × 10^−5^, L2 penalty 1.49 × 10^−6^, split penalty 3.38 × 10^−5^ and at most 10,000 boosting rounds. The final model was fit to all development samples (3,157,941 finite gene-sample rows), reserving a deterministic 10% row subset for early stopping (seed 42); the selected model stopped at round 7,616.

EPIC-Seq^11^ was run from BAM files using a Snakemake/SLURM implementation of the published workflow, with MANE GRCh38 v1.3 TSS definitions, WGS mode, no gene grouping and minimum mapping quality 30. The workflow extracted fragment-size distributions and oriented cfDNA fragmentation over the four EPIC-Seq promoter, flanking and nucleosome-depleted-region windows and generated promoter fragment entropy (PFE), nucleosome-depleted-region (NDR), 2-kb coverage and oriented-fragment matrices. PFE represented Bayesian fragment-length entropy relative to negative-control flanking regions. NDR_relative_2k was NDR coverage from 100–300-bp fragments within TSS ±150 bp divided by 0.001 plus total coverage within TSS ±1,000 bp. Per sample, PFE was gamma-CDF scaled and NDR values were divided by their median among EPIC-Seq housekeeping negative-control genes. The published pretrained WGS coefficient ensemble (EnsembleModelCoeffsWGS.rds) was restricted to combo models using normalized PFE and NDR_relative_2k; inferredGEP was the median linear predictor across ensemble members. Gene–sample pairs with non-finite EPIC-Seq output were excluded from calculations. This matched-input WGS comparison does not assess the performance of the optimized, targeted EPIC-Seq assay.

For Cancer Surfaceome^27^ and Druggable Genome Tier 1^28^ benchmarks, RNA-seq truth values > 2 in log_2_(CPM+1) space were designated active and lower values inactive. ROC curves used each method’s continuous point output; comparisons within each cohort and gene set used the common finite gene-sample intersection across non-naive methods. For uncertainty-informed Proteus, calls with |AOR| < 5 were abstained. PPV, NPV, sensitivity and specificity were calculated among retained calls, with coverage reported as the retained fraction.

Pearson’s *r* was used for same-scale comparisons of Proteus or XGBoost with RNA-seq. Spearman’s *ρ* was used for rank comparisons and methods not sharing the RNA-seq scale, including EPIC-Seq and IHC. RMSE and MAE used finite same-scale observations. Sample metrics were calculated across genes; gene metrics were calculated across samples.

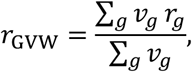

with an analogous definition for *ρ*_IJK_, where *v*_g_ is observed gene-level expression variance and *r*_g_ is gene-level correlation across samples. Genes with undefined correlation because predictions were constant or invalid were assigned zero and retained; genes with insufficient or invariant truth were excluded using a common mask.

To assess drivers of uncertainty and error, matched CRPC-WCDT and BLCA-RA gene-sample pairs were pooled. Separate crossed random-effects models were fit for 90% predictive interval width, -|AOR| and absolute residual:

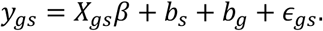

The fixed design contained cohort, tumor fraction and TSS mean depth as technical covariates and observed expression and non-PHT development-set expression variance as truth/boundary covariates; *b_s_* and *b*_g_ were sample and gene random intercepts. Models were fit using deterministic expectation-maximization with alternating best linear unbiased predictor updates for the crossed effects and generalized least-squares updates for fixed effects. Correlated technical and truth/boundary fixed-effect variance was partitioned by an order-independent two-block Shapley decomposition, then reported with sample, gene and residual variance components as fractions of total modeled variance. Gene-level diagnostics related withholding frequency (|AOR| < 5) to observed expression, distance from the activity boundary and development-set dynamic range. For the effective-coverage analysis, effective tumor coverage was defined for each eligible gene-sample pair as TFx multiplied by the gene-specific mean region fragment depth reported by Triton; low-, medium- and high-coverage groups were pooled as tertiles across eligible pairs from CRPC-WCDT and BLCA-RA.

### Gene set enrichment and PAM50 scoring

Predicted point estimates and RNA-seq expression values underwent identical pre-filtering: genes containing any missing values were dropped, and no abundance-based filtering was performed. Single-sample gene-set enrichment analysis (ssGSEA) was performed using GSEApy^73^ (v 1.1.12). Hallmark (v2026.1)^74^ gene sets were evaluated with min_size = 10 and max_size = 500. The 31-gene Cell Cycle Progression signature (CCP.31^32^) and CRPC molecular subtype-defining^33^ signatures (NE.10, ARG.10) were evaluated with min_size = 3 and max_size = 500 (**Supplementary Figs. 8b and 9; Supplementary Tables 3 and 4**).

Samples were assigned to PAM50 categories using the classification method described previously (genefu::intrinsic.cluster.predict with the pam50.scale model)^34^. We restricted the classification to Luminal A (LumA), Luminal B (LumB) and Basal, removing HER2 and Normal samples from the training set and centroid scores before classification (**Supplementary Fig. 8c– e; Supplementary Table 3**).

### Differential gene expression and pathway enrichment

Differential expression used limma-trend^75^ with two-sided moderated t statistics: lmFit, contrasts.fit, and eBayes(trend=TRUE, robust=TRUE). Proteus predictive means or RNA-seq values formed the expression matrices. Genes with any non-finite value in the aligned matrix were removed, and retained genes were required to have log₂(CPM+1) > 1 in at least three samples. Observation-level precision weights were derived from the width of the 90% predictive interval:

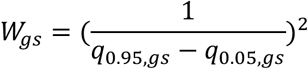

where q_0.05,gs_ and q_0.95,gs_ are the predictive 5th and 95th percentiles for gene *g* in sample *s*. Positive finite weights were normalized by their global mean to produce a mean weight of one; weights were not clipped. Genes with non-finite or non-positive aligned weights were excluded from weighted fits. Unpaired analyses used a no-intercept group-only design for the specified biological contrast. Paired pre/post-treatment analyses included patient as a blocking factor. TFx was not included as an additional covariate.

P-values were adjusted using the Benjamini-Hochberg procedure. Differentially expressed genes were defined by adjusted p < 0.05 and absolute log₂ fold change > 1. For pathway analysis, all genes with finite limma moderated t statistics were ranked and analyzed using preranked GSEA implemented in GSEApy (version 1.1.12). Hallmark v2026.1 gene sets were evaluated with min_size=10 and max_size=500; ARG.10, NE.10, CCP.31, and other custom sets used min_size=3 and max_size=500. Normalized enrichment scores and false-discovery-rate q values were reported, with FDR < 0.25 used for descriptive pathway summaries. For each gene, the abstention frequency was calculated across eligible predictions in the pooled CRPC-WCDT and BLCA-RA cohorts using the default |AOR| < 5 rule. The 1,000 genes with the highest abstention frequencies were submitted to Enrichr^76^ using the MSigDB Hallmark 2020 and Tabula Sapiens^77^ libraries. Enrichr nominal p-values were obtained from Fisher’s exact tests and adjusted within each library using the Benjamini-Hochberg procedure; complete outputs are reported in **Supplementary Table 2**.

### Statistical analyses

Unless stated otherwise, tests were two-sided and expression values were analyzed in log_2_(CPM + 1) space. Pearson (r) and Spearman (ρ) correlations and p-values used SciPy^78^ (v1.15.1). Mann-Whitney U tests compared independent groups, Wilcoxon signed-rank tests paired groups, and Kruskal-Wallis tests more than two groups. Rank-biserial or signed rank-biserial effects were reported for rank tests and Cohen’s *d* where shown. ROC curves and AUC used scikit-learn^79^ (v1.6.1). Genome-wide and multi-biomarker p-values were adjusted using the Benjamini-Hochberg procedure.

The patient was the statistical unit for repeated measurements. Paired pre/post-treatment differential expression used limma-trend with patient pairing and interval weights where indicated. Paired single-gene or pathway comparisons used Wilcoxon tests. Multiple metastatic biopsies were averaged to a patient-level transcriptome except when metastatic heterogeneity was analyzed.

Cox proportional-hazards models were fitted using Lifelines^80^ v0.30.0. For the CRPC-Pluvicto Hallmark pathway analysis, each baseline pathway score was evaluated in a separate model containing the pathway score and estimated tumor fraction as covariates. Both covariates were divided by their sample standard deviations before fitting. The reported pathway hazard ratio represents the change in the hazard of death per 1-s.d. increase in pathway score, adjusted for standardized TFx. Analyses used complete cases with available pathway score, TFx and survival data (CRPC-Pluvicto, n = 115, 96 deaths). Hazard ratios, 95% Wald confidence intervals and two-sided Wald P values were obtained for the pathway coefficient. The Figure 5b forest plot classified pathways using nominal two-sided Wald P < 0.05; BH-adjusted q values were reported in **Supplementary Table 4**. Kaplan-Meier plots used median splits and two-sided log-rank tests.

### Software and reproducibility

*Proteus* used Python 3.10, PyTorch 2.5.1 and CUDA 12.1. The canonical environment is specified in ‘proteus.env’ included in the Proteus code repository, including SciPy, scikit-learn, Zarr, Optuna, lifelines, gseapy, limma, genefu, SHAP, Captum and UMAP. Fixed seeds were used for data sampling, configured Optuna runs, attribution sampling, UMAP, bootstrap analyses and Monte Carlo draws.

### Data availability

Raw whole-genome sequencing data for CRPC-PDX plasma ctDNA are available through NCBI BioProject under accession PRJNA900550. Raw CRPC-PDX tumor RNA-seq data are available through the Gene Expression Omnibus (GEO) under accession GSE199596.

Raw genomic sequencing data for the SCLC-PDX and patient SCLC cohorts are available under controlled access through dbGaP under study accession phs003570.v1.p1.

Raw CRPC-WCDT cfDNA whole-genome and matched tumor sequencing data are available under controlled access through the European Genome-phenome Archive under study accession EGAS00001005783 and dataset accession EGAD00001008460.

Raw plasma cfDNA sequencing data from the CRPC-RA cohort are not publicly available because participant consent does not permit unrestricted genomic data sharing. Raw CRPC-RA tumor RNA-seq data are publicly available through GEO under accessions GSE147250 and GSE228283.

Raw BLCA-RA plasma cfDNA whole-genome sequencing data have been deposited under controlled access through dbGaP under study accession phs001797.v2.p1 and will be released no later than publication. Raw BLCA-RA tumor RNA-seq data are publicly available through GEO SuperSeries accession GSE302273.

Raw plasma cfDNA sequencing data from the CRPC-UW cohort are not publicly available because participant consent does not permit unrestricted genomic data sharing. Processed data originally released with that cohort are available at https://github.com/GavinHaLab/CRPCSubtypingPaper/tree/main/Data.

Raw cfDNA whole-genome sequencing data from the HD-U and HD-L cohorts will be deposited in dbGaP and made available under controlled access upon publication; the accession number will be provided upon assignment. Raw RNA-seq data from the HD-L cohort are available through GEO under accession GSE309892.

Requests for controlled access to the CRPC-RA, CRPC-UW and CRPC-Pluvicto data should be directed to Gavin Ha. Requests will receive an initial response within 30 days and will be evaluated for compatibility with participant consent and the proposed research purpose. Access requires applicable institutional and ethics approvals and execution of a data-use agreement protecting participant confidentiality. No authorship requirement is imposed as a condition of access.

All other publicly shareable processed data including processed tumor RNA-seq expression data and cfDNA-derived Proteus predictions necessary to reproduce the results are provided in Supplementary Tables 5 and 6.

### Code availability

Triton software for comprehensive cfDNA characterization is publicly available on GitHub and can be accessed at https://github.com/GavinHaLab/Triton. Proteus software will be made publicly available on GitHub prior to publication. A private, read-only link to an unpublished Zenodo record containing a complete archive of the Proteus source code, configuration files, training and optimization code, downstream analysis pipelines, trained model checkpoints, reference assets, and a single-sample demonstration with a quick-start guide and expected outputs has been supplied directly to the journal with this submission for confidential access by editors and reviewers. To preserve confidentiality, the access link is provided separately in the submission materials rather than reproduced in the manuscript. The archive is provided solely for evaluation of this manuscript and is not licensed for public use or redistribution.

## ACKNOWLEDGEMENTS

We thank the many patients and their families for their altruistic contributions to this study. We thank members of the Ha and Nelson research groups for helpful discussions. We thank Dr. Robert B. Montgomery and the Molecular Correlates Biospecimen Blood Collection and Sample Delivery Clinical Research Support Team, and the Biospecimen Processing & Biorepository Shared Resources of Fred Hutch for assistance with blood processing. We also thank the Genomics & Bioinformatics Shared Resource (RRID:SCR_022606) of the Fred Hutch/University of Washington/Seattle Children’s Cancer Consortium (P30CA015704), the Stuart and Molly Sloan Precision Oncology Institute, and the Institute for Prostate Cancer Research clinicians and staff who support the University of Washington rapid autopsy program and the LuCaP PDX program. This work was supported by the National Institutes of Health (DP2 CA280624, R01CA280056, P50CA097186, P30CA015704, P01CA163227, R50CA27433, R01CA281801), CDMRP awards (W81XWH-21-1-0513, PC230582), a Prostate Cancer Foundation Young Investigator Award (R.D.P., generously funded by the Milken family), a Mahan postdoctoral fellowship (A.P.M.) and the FHCC Scientific Computing Infrastructure award (ORIP Grant S10OD028685).

## CONTRIBUTIONS

R.D.P., P.S.N., G.H. conceived and designed the study.

R.D.P. designed, developed, and implemented Proteus and Triton methods and software.

Al.N., Ak.N., G.H. contributed to the design, development, or validation of Proteus and Triton.

R.D.P., T.W.P., I.M.C., P.C.G., P.I., P.C., M.A., M.V., A.P.M., A.P., P.S.N., G.H. performed the computational analysis, data curation, and/or data interpretation. R.D.P., Al.N. generated the data visualization.

P.C.G., J.B.H., R.D., L.K., H-M.L., C.M., D.M., A.C.H., P.S.N., M.T.S. contributed specimens or sequencing data, processed specimens, and/or clinical interpretations.

R.A.D., A.G., H-M.L., C.M., A.I., D.L.C., A.C.H., P.S.N. recruited patients, obtained patient consent, coordinated acquisition of clinical specimens, and contributed clinical data and interpretations.

E.S., M.C.H. provided pathology assessment and data interpretations.

R.D.P., P.S.N., G.H. wrote the original draft of the manuscript.

R.D.P., I.M.C., P.C.G., P.I., M.A., H-M.L., C.M., A.I., M.C.H., P.S.N., G.H. reviewed and edited the manuscript.

P.S.N., G.H. supervised the study.

R.D.P., P.S.N., G.H. acquired funding for the study.

## COMPETING INTERESTS

A patent application has been filed on methodologies developed from this work.

G.H. receives research support from Pfizer and has consulted for Quest Diagnostics; all activities are unrelated to this work.

A.I. received fees for advisory work from Curium, ITM, Novartis, Lantheus, Bayer, Boston Scientific and Ambrx/J&J (through institution). A.I. receives research funding from Novartis, SNMMI and NCI/NIH.

All other authors declare no competing interests.

## Notes

### Summary of Updates

This revision substantially expands the patient datasets and validation of Proteus and reframes the manuscript around rigorous benchmarking and uncertainty-aware inference. We added direct comparisons with EPIC-Seq and a machine learning model, including controlled evaluations across sequencing depths and tumor fractions and metrics designed to capture patient-specific variation in individual genes. Proteus was extended to produce probabilistic expression estimates, prediction intervals, and confidence measures that allow uncertain predictions to be withheld or down-weighted, and these outputs were evaluated in independent patient cohorts. The revised manuscript also includes expanded analyses of therapeutic targets, tumor phenotypes, pathway activity, and clinical applications, including larger survival analyses and new longitudinal treatment analyses. The title, text, figures, Methods, supplementary figures, and supplementary tables have been extensively updated.

