## Supplementary Materials for "Gene expression inference from cell-free DNA using uncertainty-aware deep learning"

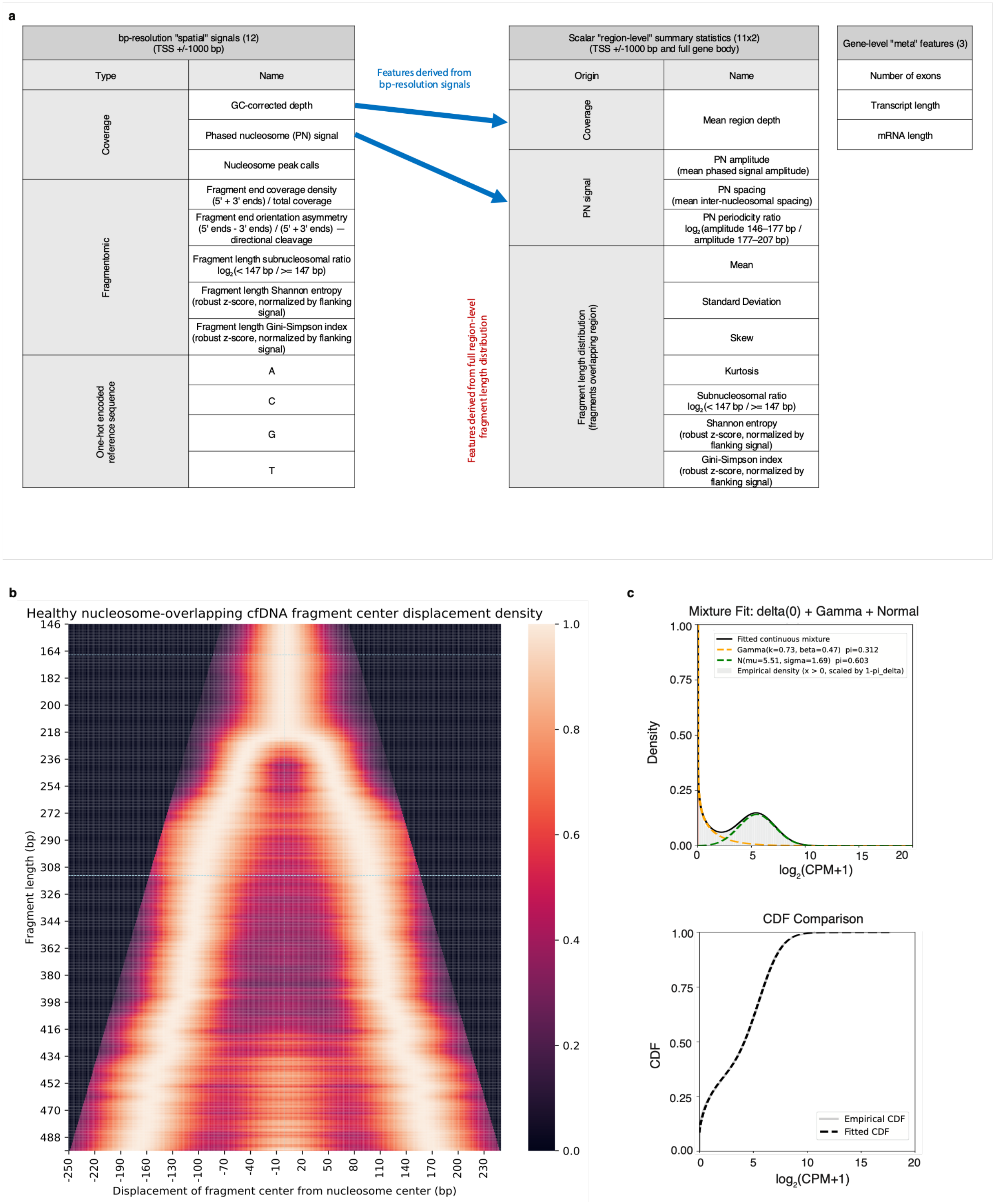


***Supplementary Figure 1. Triton feature extraction and Proteus probabilistic output.***

a) Inventory of Proteus inputs generated by Triton or obtained from reference annotation. TSS-centered inputs comprise eight biological signals across a 2,000-bp TSS window: GC-corrected depth, phased-nucleosome (PN) signal, PN peak calls, fragment-end density, fragment-end orientation asymmetry, subnucleosomal-fragment ratio, fragment-length Shannon entropy and fragment-length Gini–Simpson diversity, together with four one-hot reference-sequence channels. Eleven region-level summaries are calculated independently over the TSS window and gene body: mean depth; PN amplitude, mean spacing and periodicity ratio; and fragment-length mean, standard deviation, skewness, kurtosis, subnucleosomal ratio, Shannon entropy and Gini–Simpson diversity. Exon count, mature mRNA length and full transcript span are supplied separately.

b) Empirical displacement-density matrix used to reweight nucleosome-overlapping cfDNA fragments by fragment length and displacement of the fragment center from the inferred nucleosome center. Profiles were derived from healthy-donor cfDNA over 186 high-confidence nucleosome centers shared across reference tissues and Gaussian smoothed as described in the Methods. Dashed lines mark representative mono- and dinucleosomal fragment lengths; color denotes normalized displacement density.

c) Illustrative Proteus hurdle–Gamma–Normal output distribution derived from bulk RNA-seq training labels. The upper plot shows component densities, mixture weights and the fitted continuous mixture over positive expression; the lower plot compares empirical and fitted cumulative distributions. Expression is $\log_2(\mathrm{CPM} + 1)$.

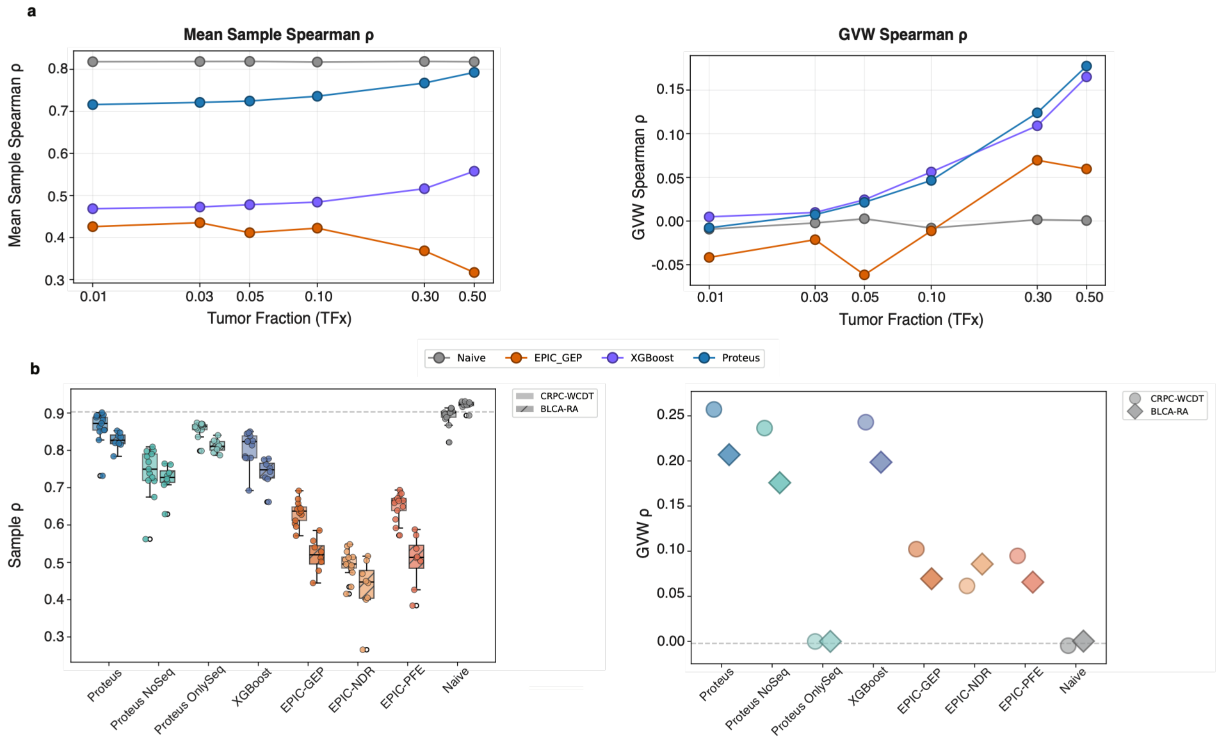


***Supplementary Figure 2. Low-depth benchmarking and input-modality ablation.***

a) Controlled benchmarking at 5× WGS depth using admixtures derived from PDX models excluded from all model development. Tumor fractions range from 0.01 to 0.50, with n = 10 admixtures per condition. Proteus is compared with XGBoost trained on Triton region-level features, EPIC-Seq inferred gene-expression profiles (EPIC-GEP) and the naïve RNA-seq control. Left, mean within-sample Spearman correlation across genes; right, gene-variance-weighted Spearman correlation across samples ($\rho_{GVW}$). Points show each TFx condition.

b) Benchmarking in patient cohorts excluded from development: external CRPC-WCDT (n = 13 patients) and cancer-type-held-out BLCA-RA (n = 8 patients). Full Proteus is compared with no-sequence and sequence-only ablations, XGBoost/Triton, EPIC-GEP, EPIC nucleosome-depleted-region (EPIC-NDR) and promoter-fragment-entropy (EPIC-PFE) scores, and the naïve control. Left, within-sample Spearman correlations across genes; points are patients, boxes show median and interquartile range, and whiskers extend to 1.5 times the interquartile range. Right, cohort-level $\rho_{GVW}$. Circles denote CRPC-WCDT and diamonds BLCA-RA; the dashed line marks $\rho_{GVW} = 0$.

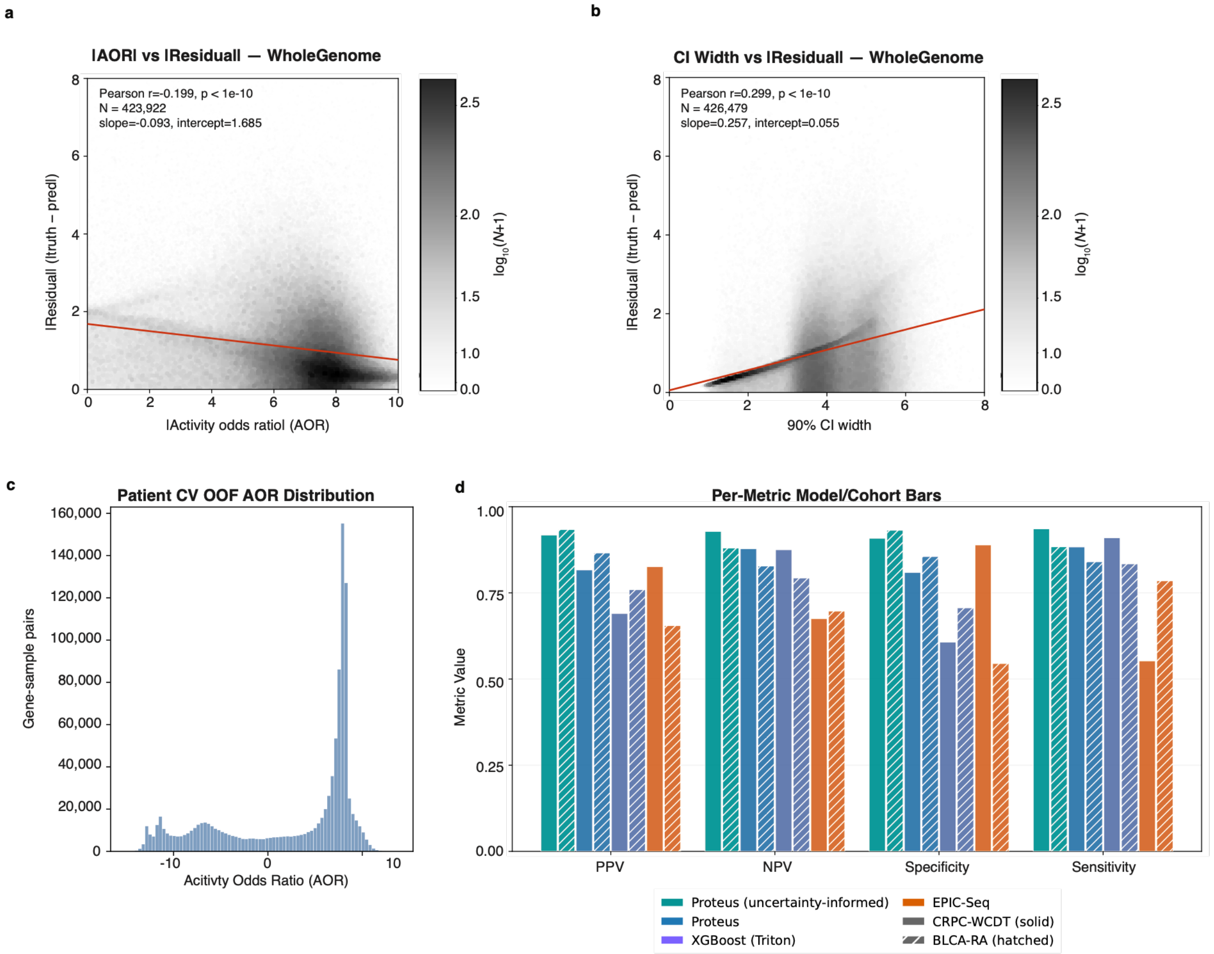


***Supplementary Figure 3. Proteus uncertainty measures identify reliable gene-activity calls.***

a,b) Absolute prediction residual versus absolute AOR (a; n = 423,922 gene–sample pairs) and central-90%-model-interval width (b; n = 426,479 gene–sample pairs) in pooled held-out CRPC-WCDT and BLCA-RA data. Gray density is $\log_{10}(N + 1)$ observations per bin; red lines are ordinary least-squares fits. Two-sided Pearson correlations and P values are displayed.

c) Distribution of total gene-sample pair AOR values in cross-validation out-of-fold results from patient samples used in training (CRPC-RA and SCLC).

d) Cancer Surfaceome activity benchmarking in CRPC-WCDT and BLCA-RA across tested models, showing PPV, NPV, specificity and sensitivity (among retained calls in the case of uncertainty-informed Proteus). Solid bars denote CRPC-WCDT and hatched bars BLCA-RA

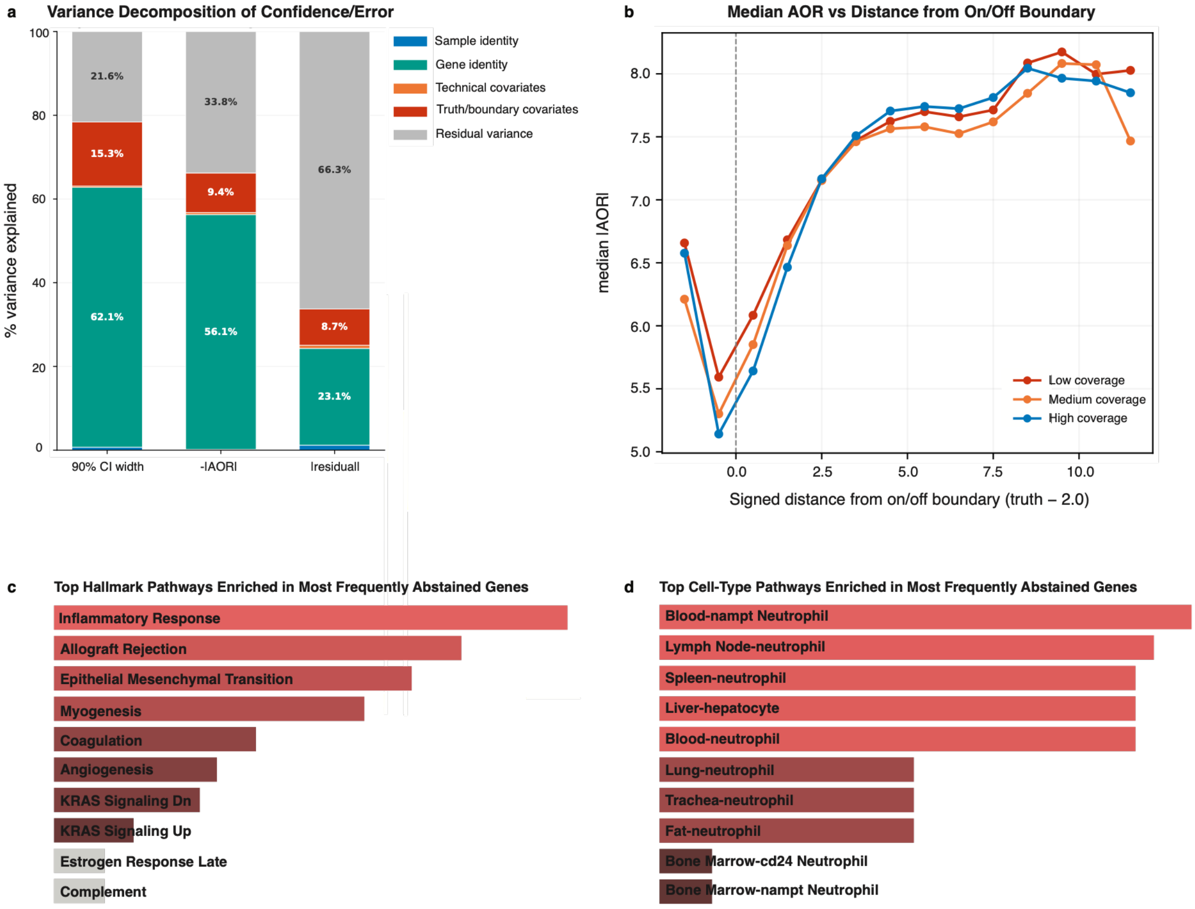


***Supplementary Figure 4. Biological and technical determinants of Proteus uncertainty and error.***

a) Variance decomposition of central-90%-model-interval width, –|AOR| and absolute residual in pooled CRPC-WCDT and BLCA-RA gene–sample pairs. Crossed random-effects models partition variance into sample, gene, technical covariates (cohort, TFx and TSS mean depth), truth/boundary covariates (observed expression and development-set expression variance) and residual. Bars show the percentage of total modeled variance assigned to each component.

b) Median |AOR| versus signed distance from the $\log_2(\text{CPM} + 1) = 2$ activity boundary, stratified by effective tumor coverage. Effective coverage is TFx multiplied by Triton gene-specific mean region depth; low, medium and high groups are pooled tertiles across eligible CRPC-WCDT and BLCA-RA pairs. The dashed line marks the activity boundary.

c,d) Top MSigDB Hallmark 2020 (c) and Tabula Sapiens cell-type (d) terms among the 1,000 genes most frequently withheld (|AOR| < 5) in pooled CRPC-WCDT and BLCA-RA predictions.

Enrichr terms are ranked by nominal Fisher exact-test P value; bar length and shading follow the default Enrichr significance encoding. In c, inflammatory response, allograft rejection, epithelial–mesenchymal transition, myogenesis and coagulation remained significant after Benjamini–Hochberg adjustment (adjusted $P < 0.05$). No Tabula Sapiens term remained significant. Complete results are in Supplementary Table 2.

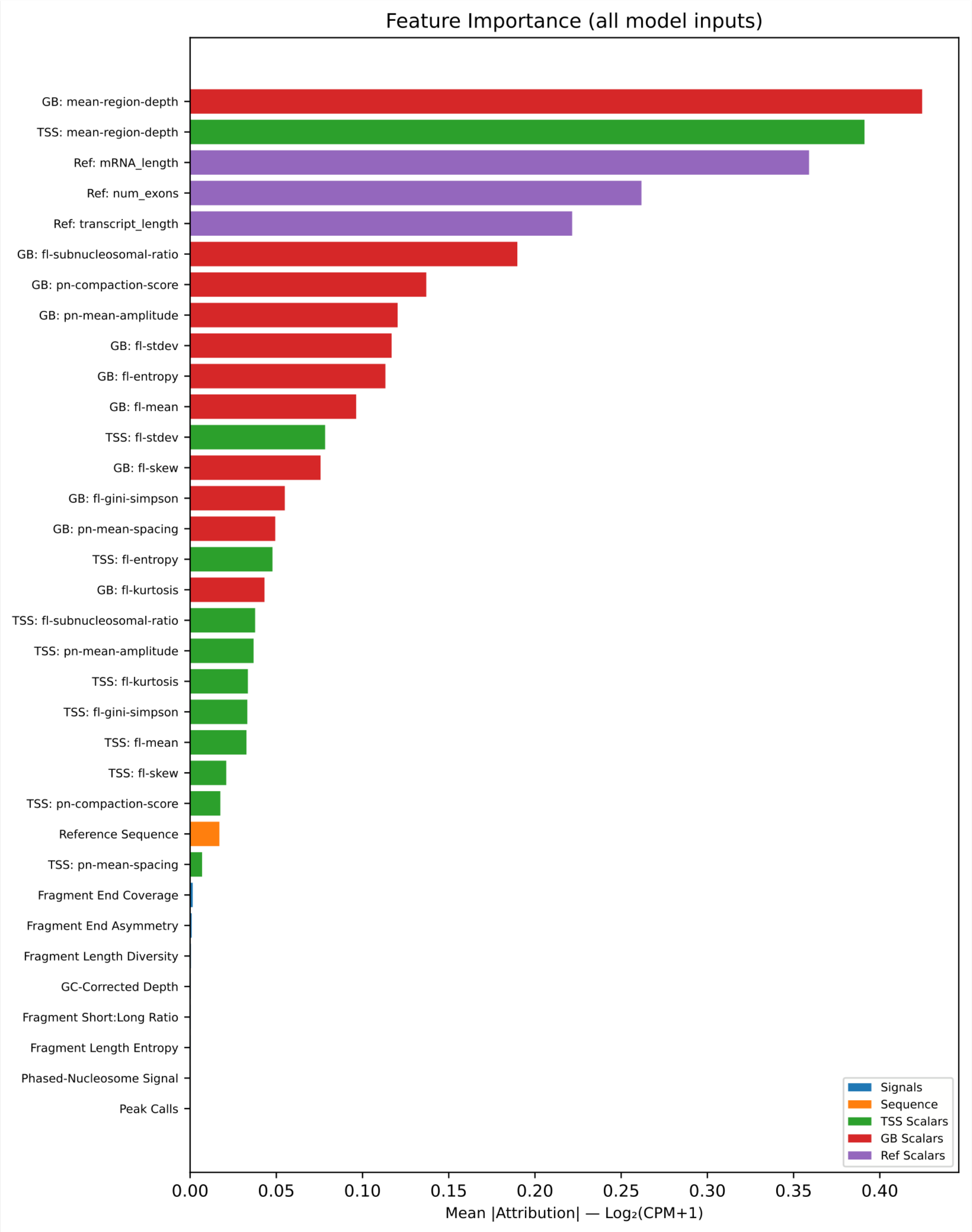


***Supplementary Figure 5. Global importance of Proteus inputs.***

Mean absolute SHAP expected-gradient attribution for all Proteus inputs, in predicted $\log_2$(CPM + 1) units and ordered by decreasing importance. Region-level features are shown separately for gene body (GB) and the TSS window; base-pair-resolution TSS channels are summarized by signal class. Colors denote spatial signal, promoter sequence, TSS region-level features, gene-body region-level features and reference annotations.

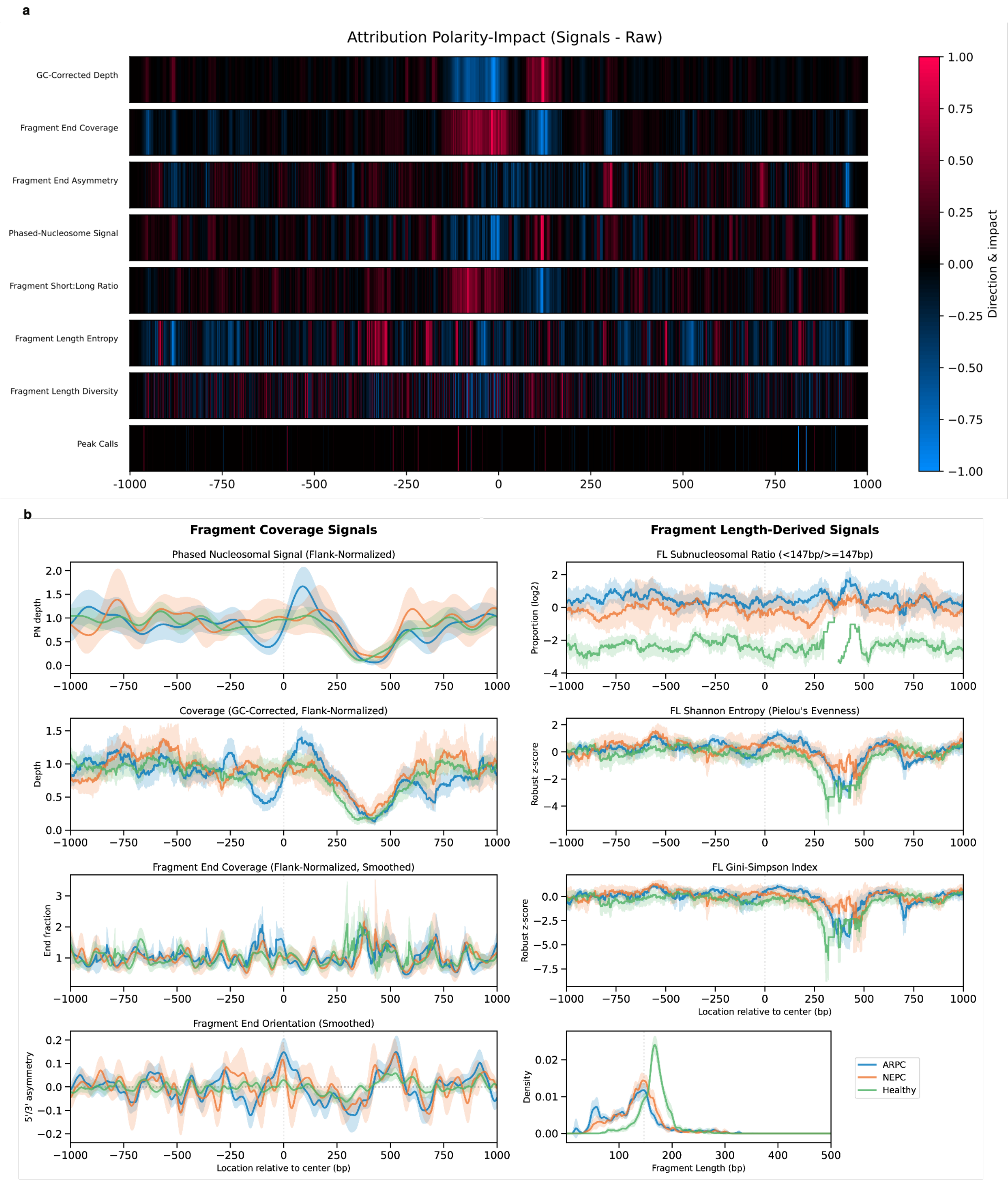

a
Attribution Polarity-Impact (Signals - Raw)
GC-Corrected Depth
Fragment End Coverage
Fragment End Asymmetry
Phased-Nucleosome Signal
Fragment Short:Long Ratio
Fragment Length Entropy
Fragment Length Diversity
Peak Calls
-1000
-750
-500
-250
0
250
500
750
1000
1.00
0.75
0.50
0.25
0.00
−0.25
−0.50
−0.75
−1.00
Direction & impact
b
Fragment Coverage Signals
Fragment Length-Derived Signals
Phased Nucleosomal Signal (Flank-Normalized)
PN depth
FL Subnucleosomal Ratio (<147bp/>=147bp)
Proportion (log2)
Coverage (GC-Corrected, Flank-Normalized)
Depth
FL Shannon Entropy (Pielou's Evenness)
Robust z-score
Fragment End Coverage (Flank-Normalized, Smoothed)
End fraction
FL Gini-Simpson Index
Robust z-score
Location relative to center (bp)
Fragment End Orientation (Smoothed)
5'/3' asymmetry
Location relative to center (bp)
Density
Fragment Length (bp)
ARPC
NEPC
Healthy

***Supplementary Figure 6. Spatial attribution patterns and representative Triton profiles.***

a) Signed SHAP polarity-impact across the 2,000-bp TSS window for the eight biological Triton channels. Red denotes positions where higher signal is associated with higher predicted expression, blue the converse and black limited directional impact. Position zero marks the TSS.

b) Representative AR TSS profiles aggregated across LuCaP ARPC PDX ctDNA, LuCaP NEPC PDX ctDNA and healthy-donor cfDNA. Coverage-related signals include PN signal, GC-corrected depth, fragment-end density and fragment-end orientation asymmetry; fragment-length signals include subnucleosomal ratio, Shannon entropy, Gini–Simpson diversity and fragment-length density. Lines show group means and bands 95% confidence intervals.

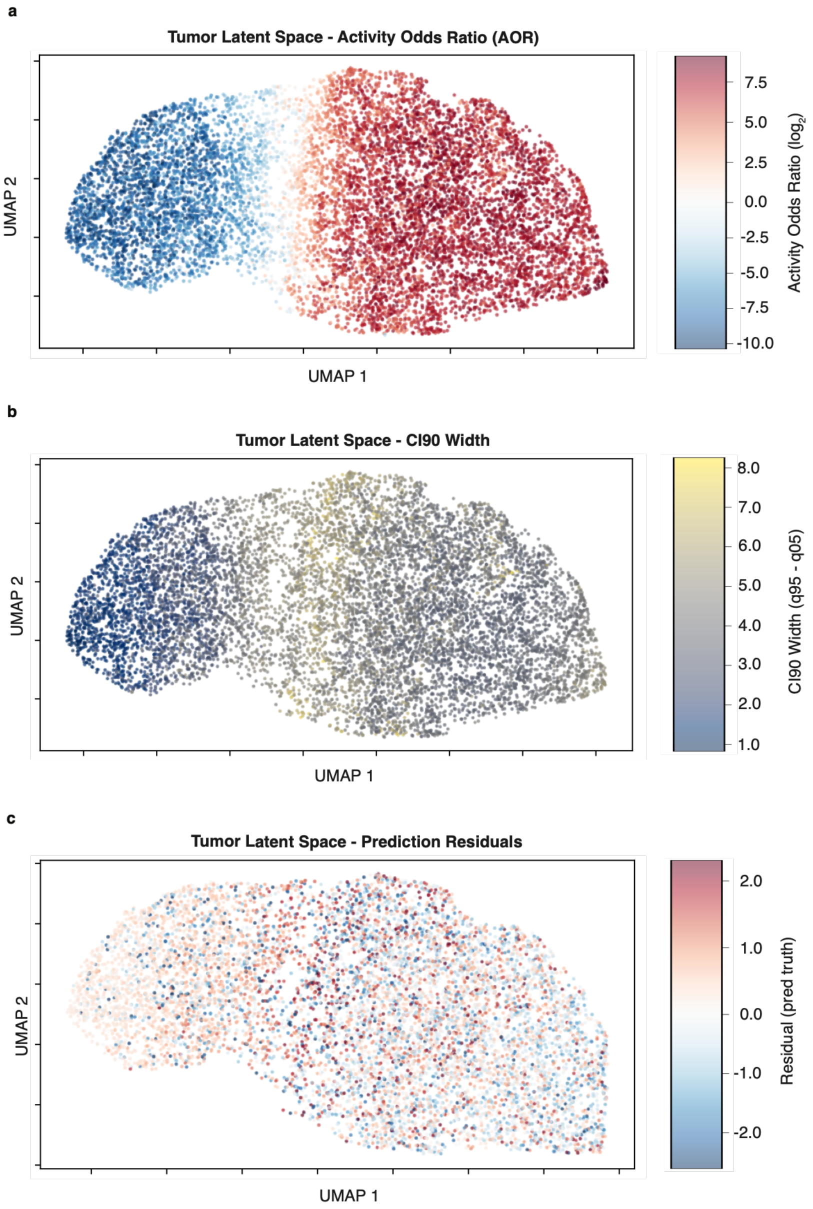


***Supplementary Figure 7. Latent-space organization of uncertainty and prediction error.***

a–c) UMAP projection of deterministic Proteus tumor-latent means for the same gene–sample examples, colored by AOR (a), central-90%-model-interval width (q0.95 – q0.05; b) or signed residual (predicted minus observed expression; c). Uncertainty and residuals concentrate near the transition between low- and high-activity regions. UMAP is used for visualization only.

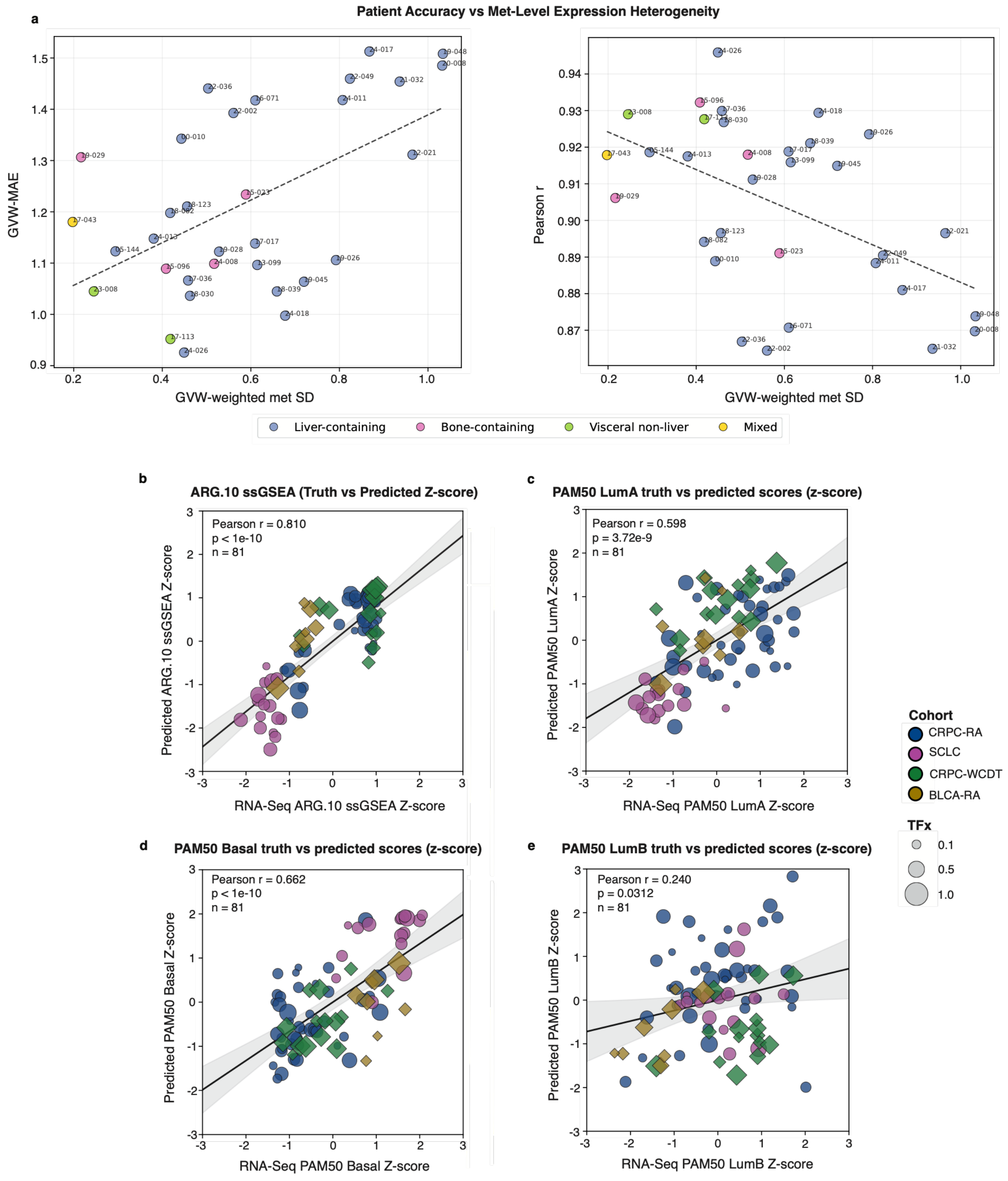

Patient Accuracy vs Met-Level Expression Heterogeneity
a
GVW-MAE
GVW-weighted met SD
Pearson r
GVW-weighted met SD
Liver-containing
Bone-containing
Visceral non-liver
Mixed
b
ARG.10 ssGSEA (Truth vs Predicted Z-score)
Pearson r = 0.810
p < 1e-10
n = 81
Predicted ARG.10 ssGSEA Z-score
RNA-Seq ARG.10 ssGSEA Z-score
c
PAM50 LumA truth vs predicted scores (z-score)
Pearson r = 0.598
p = 3.72e-9
n = 81
Predicted PAM50 LumA Z-score
RNA-Seq PAM50 LumA Z-score
d
PAM50 Basal truth vs predicted scores (z-score)
Pearson r = 0.662
p < 1e-10
n = 81
Predicted PAM50 Basal Z-score
RNA-Seq PAM50 Basal Z-score
e
PAM50 LumB truth vs predicted scores (z-score)
Pearson r = 0.240
p = 0.0312
n = 81
Predicted PAM50 LumB Z-score
RNA-Seq PAM50 LumB Z-score
Cohort
CRPC-RA
SCLC
CRPC-WCDT
BLCA-RA
TFx
0.1
0.5
1.0

***Supplementary Figure 8. Metastatic heterogeneity and recovery of gene-set scores.***

a) Within-patient metastatic expression heterogeneity versus Proteus accuracy in CRPC-RA patients with multiple sampled metastases (n = 32 patients). Heterogeneity is gene-variance-weighted RNA-seq standard deviation across lesions. Accuracy is gene-variance-weighted MAE (left) or Pearson correlation between patient-mean RNA-seq and Proteus prediction (right). Points are patients and colors denote lesion-site distribution; dashed lines are linear fits. Two-sided Spearman $\rho = 0.43$, $P = 0.013$ for error and $\rho = –0.46$, $P = 0.009$ for correlation.

b) RNA-seq-derived versus Proteus-derived ARG.10 ssGSEA z-scores across matched cohorts (n = 81 patients; two-sided Pearson r = 0.810, P < 1 × 10–10).

c–e) RNA-seq-derived versus Proteus-derived PAM50 centroid z-scores for Luminal A (c; r = 0.598, P = 3.72 × 10–9), Basal (d; r = 0.662, P < 1 × 10–10) and Luminal B (e; r = 0.240, P = 0.0312) across matched cohorts (n = 81 patients; two-sided Pearson tests). In b–e, color denotes cohort, circles out-of-fold predictions, diamonds held-out predictions, point size TFx, lines ordinary least-squares fits and bands 95% confidence intervals around the fitted mean.

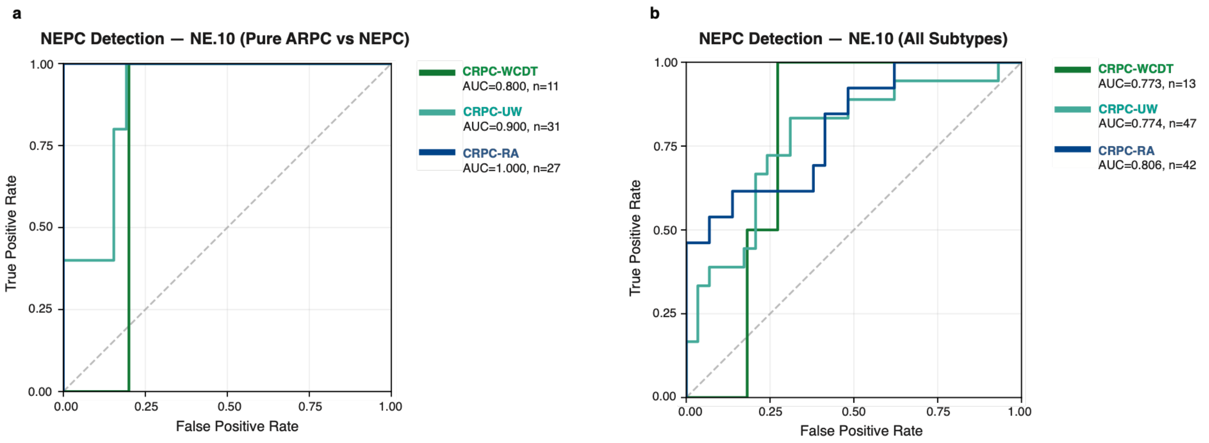


***Supplementary Figure 9. Neuroendocrine prostate cancer classification using Proteus-derived NE.10 scores.***

a) ROC curves for pure ARPC versus pure NEPC classification using Proteus-derived NE.10 scores in CRPC-WCDT (n = 11 cfDNA samples; AUC = 0.800), CRPC-UW (n = 31 samples; AUC = 0.900) and CRPC-RA (n = 27 patients; AUC = 1.000).

b) ROC curves for detection of any neuroendocrine component across annotated subtypes in CRPC-WCDT (n = 13 samples; AUC = 0.773), CRPC-UW (n = 47 samples from 27 patients; AUC = 0.774) and CRPC-RA (n = 42 patients; AUC = 0.806). Dashed diagonals denote chance.

### SUPPLEMENTARY TABLE AND DATA LEGENDS

***Supplementary Table 1. Cohort, sample and model-development metadata.***

Sample-level metadata for all cohorts appearing in this manuscript (PDX, healthy-donor and patient cohorts), including sample identifier, cohort, collection time point where applicable, WGS depth, tumor fraction estimates, clinical phenotype if available, survival metrics where applicable, and analytical use (development, cross-validation, held-out evaluation or clinical application). Development samples include fixed five-fold assignments. In silico admixture records additionally report PDX and healthy-donor parents, target TFx and target depth, and designation as training (PHT) or benchmarking (PHS) mixtures.

***Supplementary Table 2. Benchmarking and uncertainty-analysis results.***

Complete performance results for Proteus, Proteus input ablations, XGBoost/Triton, EPIC-Seq and benchmark controls across in silico admixtures and held-out CRPC-WCDT and BLCA-RA cohorts. Reported outputs include sample-level and gene-level correlations, gene-variance-weighted correlations, error metrics, call coverage, PPV, NPV, sensitivity and specificity for Cancer Surfaceome and Druggable Genome Tier 1 sets. Additional worksheets contain variance-component estimates and the complete Enrichr outputs for the 1,000 most frequently withheld genes against the MSigDB Hallmark 2020 and Tabula Sapiens libraries, including term, overlap, nominal P value, Benjamini-Hochberg-adjusted P value, odds ratio, combined score and overlapping genes. Also included are sample-level accuracy metrics between RNA-seq and predictive point estimates at the transcriptome level, by cohort: CRPC-RA and SCLC use out-of-fold cross-validation predictions while CRPC-WCDT and BLCA-RA are held-out predictions.

***Supplementary Table 3. Matched-cohort molecular-phenotype and orthogonal-validation results.***

Patient-level and summary results for analyses using matched cfDNA and tumor measurements in CRPC-RA, SCLC, CRPC-WCDT and BLCA-RA. Worksheets include metastatic-heterogeneity metrics, RNA-seq- and Proteus-derived ssGSEA and PAM50 scores, and marker-level comparisons among Proteus predictions, tumor RNA-seq and IHC. Exact sample counts, correlation coefficients, two-sided P values and multiple-testing-adjusted values are reported for each comparison.

***Supplementary Table 4. Clinical-cohort molecular scores and statistical analyses.***

Results for clinical application cohorts without contemporaneous matched tumor RNA-seq. The CRPC-UW worksheet contains pathway enrichment scores based on Proteus predictions. CRPC-Pluvicto worksheets contain baseline Hallmark scores; TFx-adjusted Cox model coefficients, hazard ratios, 95% Wald confidence intervals, two-sided Wald P values and Benjamini-Hochberg q values for Hallmark pathways; paired pre/post-treatment limma results with and without predictive-interval weights; and corresponding preranked GSEA results.

***Supplementary Table 5. Processed RNA-seq expression values used for model development and evaluation.***

TMM-normalized tumor or blood RNA-seq expression values in $\log_2$(CPM + 1) space for all samples with expression labels, organized by cohort and indexed by MANE Select v1.3 gene identifier and sample identifier. Where multiple metastatic biopsies were available (CRPC-RA), both lesion-level values and the patient-level mean used for model training or evaluation are provided and explicitly labeled.

***Supplementary Table 6. Proteus-inferred and comparator expression outputs for all analyzed cfDNA samples.***

Proteus predictive means for each analyzed gene–sample pair, indexed by MANE Select v1.3 gene and sample identifiers. Cohorts include development and benchmarking admixtures, matched patient cohorts and clinical application cohorts. Both XGBoost and EPIC-Seq predictions are also included for those cohorts used in direct benchmarking (PHS, CRPC-WCDT, and BLCA-RA).
